# Metformin-Induced Mitochondrial Remodeling Creates Adaptive and Targetable Vulnerabilities in Cancer Cells

**DOI:** 10.1101/2025.05.29.656565

**Authors:** Firoz Khan Bhati, Uday Saha, Sandeep Singh, Manoj Kumar Bhat

## Abstract

Mitochondrial remodeling has emerged as a key regulator of cancer cell metabolism, cell stemness, development of drug resistance, and induction of apoptosis. Metformin, a clinically approved inhibitor of mitochondrial complex I, has shown variable antitumor activity in preclinical and clinical settings, implying that adaptive mitochondrial responses can buffer its cytotoxic effects. Here, we show that metformin reprograms mitochondrial dynamics in liver and colorectal cancer cells, increases mitochondrial biomass, alters fusion–fission balance, and enhances biogenesis in a cell type–dependent manner. These adaptations preserve mitochondrial function and support survival under metformin-induced energetic stress. Notably, the targeted disruption of these mitochondrial adaptations induces cell death in metformin-treated cancer cells and suppresses tumor growth. Overall, the study identifies metformin-driven mitochondrial remodeling as a central adaptive axis to complex I inhibition and provides a rationale for co-targeting mitochondrial dynamics and metabolism to enhance the anticancer efficacy of metformin.

## Introduction

Mitochondria undergo fission, fusion, biogenesis, and mitophagy to maintain their energetic and structural integrity. These interconnected processes, collectively referred to as mitochondrial dynamics, regulate cellular metabolic capacity. In cancer cells, mitochondrial remodeling supports energy production, biosynthetic demands, and buffers metabolic stress induced by nutrient deprivation, hypoxia, or therapeutic interventions (Giannitti *et al*, 2025; McGuirk *et al*, 2020; Tufail *et al*, 2024; Chiche *et al*, 2010; Andrzejewski *et al*, 2018). Tumors frequently exploit these alterations to facilitate survival and metastasis. Several therapeutic agents including biguanides, rotenone derivatives, and DHODH inhibitors directly disrupt mitochondrial oxidative metabolism. Among these, metformin, an FDA-approved antidiabetic drug, has gained attention as a mild and clinically tolerable inhibitor of mitochondrial complex I. Nevertheless, its anticancer efficacy remains inconsistent across clinical settings, suggesting the involvement of adaptive metabolic reprogramming that limits therapeutic benefit(Mukherjee *et al*, 2023; Urra *et al*, 2017; Masoud *et al*, 2020; Barakat *et al*, 2022; Lord & Harris, 2023). Mitochondrial complex I, a core component of the electron transport chain, plays a crucial role in cancer cell proliferation, resistance to apoptosis, and metastasis. Although the therapeutic targeting of complex I has been extensively investigated, strong inhibitors have not advanced further clinically due to toxic effect. This limitation has renewed interest in mild inhibitors like metformin, which is generally well-tolerated and exhibits minimal drug-associated toxicity (Urra *et al*, 2017; Masoud *et al*, 2020; Zhou *et al*, 2023; Wheaton *et al*, 2014; Vasan *et al*, 2020; Alghandour *et al*, 2021; Barakat *et al*, 2022). Additionally, metformin’s potential is being explored in combination with chemotherapy, radiotherapy, and immunotherapy (Lord & Harris, 2023). Despite the documented link between metformin-induced energy stress and adaptive mitochondrial remodeling, the functional significance of these adaptive responses in cancer cell survival remains poorly understood. Moreover, whether disrupting these adaptations can enhance metformin’s therapeutic efficacy and overcome resistance is unclear(Andrzejewski *et al*, 2018; Alamri *et al*, 2021; Scatena *et al*, 2018; Wang *et al*, 2019; Su *et al*, 2023; Ma *et al*, 2020; Lord *et al*, 2018).

In this study, we demonstrate that metformin induces mitochondrial remodeling as a stress-adaptive survival mechanism in hepatocellular carcinoma cells. We also show that pharmacologic inhibition of mitochondrial biogenesis (via doxycycline) or DHODH (via leflunomide) unmasks a synthetic lethal vulnerability, resulting in enhanced apoptosis and suppressed tumor growth in vivo. Our findings position mitochondrial remodeling as a central adaptive mechanism to complex I inhibition and highlight its co-targeting as a promising strategy to potentiate metformin’s anticancer activity.

## Materials and Methods

### Cell culture and cell lines

Human hepatocellular carcinoma cell line Huh-7 was kindly provided by Dr. Ralf Bartenschlager (University of Heidelberg, Germany). MC-38 (murine colon adenocarcinoma) was gifted by Dr. Ruben Hernandez (CIMA, Spain). 4T1 tdTomato positive (murine triple-negative breast cancer) cell line was obtained from the late Dr. Mohan Wani (BRIC-NCCS, Pune, India), originally procured from Imperial Life Science (P) Ltd. Other cell lines including B16F10 (murine melanoma), Hep-3B, Hep-G2 (human hepatocellular carcinoma), HCT116 (human colon adenocarcinoma), HT-29, and HCT15 (human colorectal carcinoma) were acquired from the American Type Culture Collection (ATCC, Manassas, VA, USA) and maintained at BRIC-NCCS’s National Cell Repository. All cell lines were tested negative for mycoplasma contamination, and human cell lines were authenticated in-house by STR analysis. Cells were cultured in Dulbecco’s Modified Eagle’s Medium (DMEM) supplemented with 10% fetal bovine serum (FBS; Gibco, NY, USA), 100 µg/mL streptomycin, and 100 U/mL penicillin (Invitrogen Life Technologies, CA, USA), under controlled conditions (37 °C, 5% CO2) in a humidified incubator (Thermo Fisher Scientific, OH, USA).

### Antibodies and chemicals

Primary antibodies against Actin (1:1000, Cat # Sc-1615), Vinculin (1:1000, Cat # sc73614), LDH-A (Cat # sc-137244), TFAM (Cat # sc-376672), and TOM-20 Alexa 488 (1:200, Cat # sc-17764 AF488) were purchased from Santa Cruz Biotechnology (CA, USA). HRP-conjugated secondary antibodies were also from Santa Cruz. CD-45 APCeFluor™ 780 (1:200, Cat # 30-F11) and eBioscience Foxp3 Staining perm/fix Buffer Set (Cat # 00-5523-00) were obtained from Invitrogen Life Technologies (CA, USA). Metformin (Cat # 151691) was from MP Biomedicals LLC (OH, USA). Doxycycline (Cat # D9891), Propidium Iodide (Cat # p-4170), MTT (Cat # USB-19265), and Crystal Violet (Cat # C-3886) were from Sigma (MO, USA). Leflunomide (Cat # HY-B0083), Mdivi-1 (Cat # HY-15886), and Liensinine (Cat # HY-N0484) were purchased from MedChemExpress (NJ, USA). Mito Tracker Deep Red FM (Cat # M22426) was from Invitrogen (OR, USA). FBS was sourced from Gibco (NY, USA), and penicillin/streptomycin from Invitrogen Life Technologies (CA, USA).

### MTT assay

Cells (3 × 10³ per well) were seeded in 96-well plates and incubated for 24 h. Drugs were administered according to experimental design. After treatment, media was removed, and 50 µL of 1 mg/mL MTT in media without FBS was added per well. Cells were incubated for 3 h, MTT removed, and 100 µL isopropanol added to solubilize formazan crystals, incubated for 10 min at room temperature. Absorbance was measured at 570 nm.

### Crystal violet cell biomass assay

Cells were seeded in 12- or 24-well plates and grown for 24 h. After 48 h drug treatment, cells were washed thrice with ice-cold PBS. Cells were fixed with 3% PFA for 20 min in the dark at room temperature, washed thrice with PBS, and stained with 0.05% crystal violet in PBS for 1 h. Stained cells were washed, imaged using an Olympus E-330 DSLR camera, and quantified with ImageJ software as previously described by Mayengbam *et al*, 2023).

### Cell death assay

#### Propidium Iodide (PI) exclusion assay

Cells were seeded in 12-well plates, treated as required, tripsinized, washed with PBS, and resuspended in 250 µL PBS. Before flow cytometry analysis (BD FACSCanto II, PE channel), cells were stained with 10 µg/mL Propidium Iodide for 15 min at room temperature. Ten thousand events were acquired and analyzed with FlowJo software (company).

#### Annexin V and PI Assay

Post-treatment, cells were tripsinized, washed, and resuspended in 1x binding buffer at 1×10⁶ cells/mL. Cells were incubated with 0.2 µL Annexin V antibody per 50 µL in binding buffer for 15 min, then stained with 10 µg/mL PI for a further 15 min. After adding 200 µL binding buffer, cells were analyzed by FACS. Total apoptosis was calculated by summing Annexin V positive, PI positive, and dual-positive cells.

### RNA extraction

After treatment, cells were washed with PBS, lysed in 1 mL Trizol, and incubated for 5 min. Chloroform (0.2 mL per mL Trizol) was added; samples were incubated 3 min and centrifuged (12,000 g, 4 °C, 15 min). The aqueous phase was collected, mixed with 0.5 mL isopropanol, incubated 10 min at room temperature, and centrifuged (12,000 g, 4 °C, 10 min). Pellets were washed in 75% ethanol, centrifuged (7,500 g, 4 °C, 5 min), air-dried, and resuspended in 25 µL RNAase-free water with 0.1 mM EDTA. RNA was solubilized at 55 °C for 10 min. Concentrations and purity (OD260/280) were measured by Nanodrop (Thermo Fisher Scientific, USA).

### cDNA preparation

cDNA synthesis used 2.5–5 µg RNA with oligo(dT) primers, dNTPs, RNAse OUT, first strand buffer, and 200 units Moloney Murine Leukemia Virus Reverse Transcriptase (MMLV-RT) following manufacturer protocols. cDNA was stored at −20 °C or used immediately for quantitative polymerase chain reaction (PCR) for detection of gene expression.

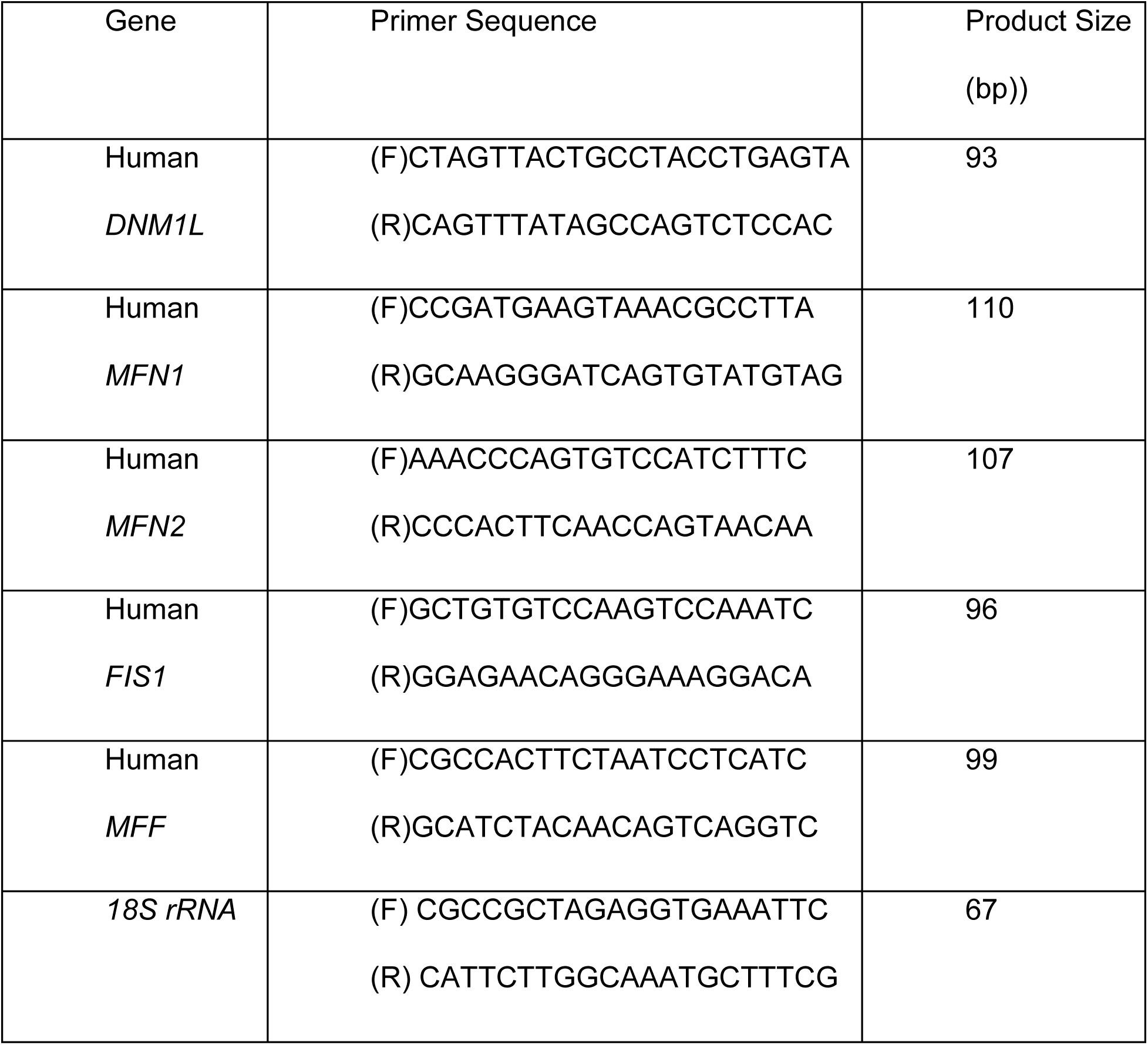

### Mitochondrial density measurement

Using Mito Tracker Deep Red FM: Cell seeded and incubated 24 h were treated as required, washed with warm DMEM (without FBS), and stained with 100 nM Mito Tracker Deep Red FM for 30 min at 37 °C. Cells were washed twice with DMEM (no FBS) and once with PBS, tripsinized and resuspended in 250 µL PBS. Data were acquired by flow cytometer and median fluorescence intensity was analyzed.

Using TOM20 antibody: After treatment, cells were tripsinized, washed with PBS, fixed with 4% PFA for 10 min at room temperature, permeabilized in chilled methanol on ice for 10 min, and washed with ice-cold PBS. Pellets were incubated with TOM20 antibody for 1 h on ice with intermittent mixing, washed, resuspended in 0.1% PFA, and analyzed by flow cytometer for median fluorescence intensity.

### Mitochondrial morphology analysis

Cells (1 × 10⁴) were seeded on coverslips in 24-well plates and treated as required. Cells were stained with 100 nM Mito Tracker Deep Red FM for 30 min at 37 °C, washed, fixed, and in the case of TOM20 staining, permeabilized with Perm buffer for 30 min. Cells were incubated with TOM20 antibody for 1 h under gentle rocking, washed thrice with PBS, and mounted with DAPI-containing medium. Imaging was performed using an Olympus FV3000 confocal microscope. Mitochondrial morphology was quantified using the mitochondrial analyzer plugin in ImageJ (Fiji) (A *et al*, 2020).

### LysoSensor Yellow-Blue assay

Hep-3B cells on coverslips were treated with DMSO (control), Mdivi-1 (20 µM), metformin

(10 mM), or their combination for 36 h, followed by incubation with 10 µM LysoSensor for 5 min. Cells were washed thrice with ice-cold PBS, fixed with 4% PFA, and imaged with excitation at 371–405 nm and emission at 420–650 nm on the Olympus FV3000 confocal microscope. Fluorescence intensities at 440 nm (blue, less acidic) and 540 nm (yellow, more acidic) were quantified and normalized to controls.

### Glucose and lactate estimation in spent media

Hep-3B cells were seeded in appropriate culture plates and allowed to attach for 24 h. Cells were then treated with metformin and the indicated mitochondrial modulators for the specified time points. At the end of treatment, culture supernatants were collected, centrifuged at 500 g for 5 min to remove debris, and the clarified media were transferred to fresh tubes. Samples were diluted as required, and glucose and lactate concentrations in the spent media were measured using Spinreact glucose and lactate assay kits (Cat. #41,013 and #1,001,330; Girona, Spain) according to the manufacturer’s instructions.

### Animal experiments

Female Balb/c and male NOD/SCID mice were procured from the in-house Experimental Animal Facility (EAF) of BRIC-NCCS, Pune, India. All the mice were housed and maintained in the animal quarters under controlled environmental conditions of 22 ± 2 °C with a light and dark cycle of 12 h, along with the free access to clean drinking water and standard rodent pellet food *ad libitum*, unless otherwise mentioned.

#### Isograft tumor model

Isografts were developed by injecting 3×10^5^ 4T1 cells in the mammary fat pad of 6-8 weeks old female BALB/c mice. A measurable tumor developed after 7 days following injection of cells in animals. Mice were randomized in two groups. Mice in group one was treated with 250 mg/kg/day metformin in drinking water whereas mice in other groups were used as control. Metformin in drinking water was replaced every 3^rd^ day as previously described (Wheaton *et al*, 2014). Mice were treated with metformin for 18 days.

Xenograft tumor model: Xenografts were developed by subcutaneously injecting 5×10^6^ Hep-3B cells in 6-8 weeks old male NOD/SCID mice. Experiment was performed twice, in one set treatment of metformin was given and, in the second set metformin, doxycycline (DXY) and leflunomide (LEF) were given as a single drug or in combination with metformin. A measurable tumor was developed after 22 days of cell injection. Before administration of drugs, mice bearing tumors ≈50-100 mm3 were randomized into required groups. Doxycycline (50 mg/kg/day) and leflunomide (7.5 mg/kg/day) were administered intraperitoneally. Doxycycline was dissolved in saline whereas leflunomide was initially dissolved in DMSO and further resuspended in corn oil which resulted in a mixture of 5%DMSO and 95% corn oil. Vehicles of these drugs were administered in control groups.

Tumors were measured by using a digital vernier caliper at 3 days intervals. Tumor volumes were calculated by using the formula: 0.523 x (Length) x (Width)^2^. When the tumor size reached around 50-100 mm^3^, tumor bearing mice were randomly distributed into control and 5 experimental groups, each containing 4 to 6 animals (Experimental group-1; Metformin, group-2; Leflunomide, group-3; Metformin and Leflunomide, group4; Doxycycline, group-5; Metformin and Doxycycline). After completion of the treatment period mice were sacrificed and tumors were excised. Tumor pictures were taken. Tumor tissues were further processed for the preparation of single cell suspension for the FACS analysis. Part of single cell suspension was utilized for western blot analysis and RT-PCR.

### RNAseq data analysis

Raw Bulk RNA sequence count data of metformin treated cancer cells or primary cells from normal individuals treated with metformin was downloaded from GEO datasets (GSE190076, GSE208245, GSE181521, GSE137317, GSE153792). Non-confirmed or unmapped reads were removed prior to analysis. Genes having low counts i.e. less than 5 in at least 60% of the samples were not considered. R (R version 4.4.0) package edgeR (Version 4.2.1) was used to normalize the counts by the TMM method to obtain CPM values. To compare the treatment groups, CPM values were log-transformed and data was plotted.

### Hematoxylin and eosin staining

Organs (heart, kidney, lung, liver) and tumor tissue were excised from the mice and stored in 10% PFA. The preserved samples were outsourced for processing for histopathological analysis (Chaitanya Laboratories, Pune, India). After processing, samples were embedded in paraffin blocks and sectioning was done. Tissue sections were deparaffinized and hydrated with xylene and graded alcohol. These were stained with hematoxylin to highlight nuclei followed by rinsing with alcohol. Next, cytoplasmic structures were stained with eosin. After further rinsing and dehydration with alcohol, the coverslips were mounted on the slides and allowed to dry. Slides were observed under a microscope and images were taken for detailed histologic / pathological analysis. The images were analyzed by pathologist and toxicity grading score was calculated as; No Abnormality Detected (NAD), Minimal changes (+1), Mild changes (+2), Moderate changes (+3), Severe changes (+4).

### Mito stress assay for mitochondrial function

Cells (1.5 × 10^4^ cells/well) were plated into 24-well XF culture plates (Cat # 100,777–004; Agilent Technologies, CA, USA) and allowed to grow overnight in a 5% CO2 incubator at 37 °C. Cells were then treated with vehicle or metformin and Mdivi-1 as a single drug or in combination for 24 h in 10% FBS-containing media and assays were performed. Cells were washed with assay medium and incubated in a humidified non-CO2 incubator at 37 °C for 45 min with 500 µl of assay medium (XF base or DMEM medium). Experiments were performed in an Agilent XF base media as per protocol provided by the Agilent Seahorse XF Cell Mito Stress Test Kit user guide. Oligomycin, carbonyl cyanide4(trifluoromethoxy) phenylhydrazone (FCCP), and rotenone & antimycin were injected through port A, B, and C respectively in each well. Oxygen consumption rate (OCR) and extracellular acidification rate (ECAR) was measured by using Seahorse XFe24 analyzer (Agilent, CA, USA).

### Statistical analysis

Statistical analysis was performed using GraphPad Prism 8. Data distribution was assessed using the Shapiro–Wilk test. For parametric data, Student’s t-test (two-tailed) was used for pairwise comparisons. Non-parametric data were analyzed using the Mann– Whitney U test. Multiple group comparisons were corrected using Tukey’s post-hoc tests following one-way or two-way ANOVA. A p-value of less than 0.05 was considered statistically significant. In all the experiments p-values is denoted as follows; *p < 0.05, **p < 0.01, ***p < 0.001, and ****p < 0.0001

## Results

### Metformin alters mitochondrial dynamics in cancer cells

Several reports have established that alterations in mitochondrial dynamics, including biogenesis, fission, fusion and mitophagy, play critical roles in sustaining cancer cell survival under stress conditions such as nutrient deprivation, hypoxia, and drug exposure (Ma *et al*, 2020). Fusion and fission are fundamental for maintaining mitochondrial homeostasis, enabling efficient energy production and adaptation to cellular stress. Fusion, mediated by proteins such as Mitofusin 1 (MFN1) and Mitofusin 2 (MFN2), allows mitochondria to merge and share contents, preserving mitochondrial integrity. Fission, regulated primarily by Dynamin-related protein 1 (DRP1) and facilitated by proteins like FIS1 and MFF, promotes mitochondrial division and isolation of damaged regions for degradation (Tábara *et al*, 2024). These dynamic processes are essential for mitochondrial responses to stress which support cancer cell survival under metabolic constraints. Previously, our group reported that energy stress promotes mitochondrial biogenesis in cancer cells (Chaube *et al*, 2015; Chaube & Bhat, 2016). Since metformin inhibits mitochondrial complex I and enhances energy stress, we investigated its effects on mitochondrial dynamics in hepatocellular carcinoma (HCC) and colorectal cancer (CRC) cells.

Mitochondrial biomass was assessed by staining with MitoTracker Deep Red FM and measuring median fluorescence intensity (MFI) via flow cytometry after 48 h treatment. Metformin significantly increased mitochondrial dye accumulation in Hep-3B, Huh-7, HCT116, HT-29, and HCT15 cells, while no measurable change was detected in Hep-G2 cells (Figure 1A). Consistent with these observations, mitochondrial biomass assessed by TOM20 staining in Hep-3B cells confirmed increased mitochondrial content following metformin treatment by confocal microscopy and flow cytometry (Figure 1B and C). Although increased lactate production has been reported to enhance mitochondrial biomass, no significant increase was detected in HCC cells treated with 10 mM lactate compared to controls (Supplementary Figure S1A-C). Confocal microscopy analysis revealed metformin-induced changes in mitochondrial morphology in Hep-3B and Huh-7 cells after 12 or 24 h, including increased circularity and reduced perimeter and area of individual mitochondria, consistent with enhanced fission (Supplementary Figure S2A-H). Western blot analysis of cells treated for 24 h showed upregulation of mt-TFA/TFAM in Hep-3B cells and decreased expression of the lower molecular weight MFF isoform in Hep-G2 cells and MFN1 expression in Huh-7 cells (Figure 1E and F). No significant changes in DRP1, the higher molecular weight MFF isoform, or MFN2 were observed across Hep-3B, Hep-G2, and Huh-7 cells (Figure 1D-F). Consistent with these protein changes, qPCR analysis revealed that metformin significantly increased the expression of the fission mediators MFF1, FIS1, and DRP1 in Hep 3B cells at 24h time point. Also the levels of MFN1 and MFN2 transcripts was increased however, the magnitude of MFN2 induction in Hep 3B, and of both MFN1 and MFN2 in Huh 7 cells was comparatively modest (Supplementary Figure S3 A-D). This imbalance suggests metformin preferentially strengthens fission machinery while only weakly engaging fusion components.

**Figure 1.**
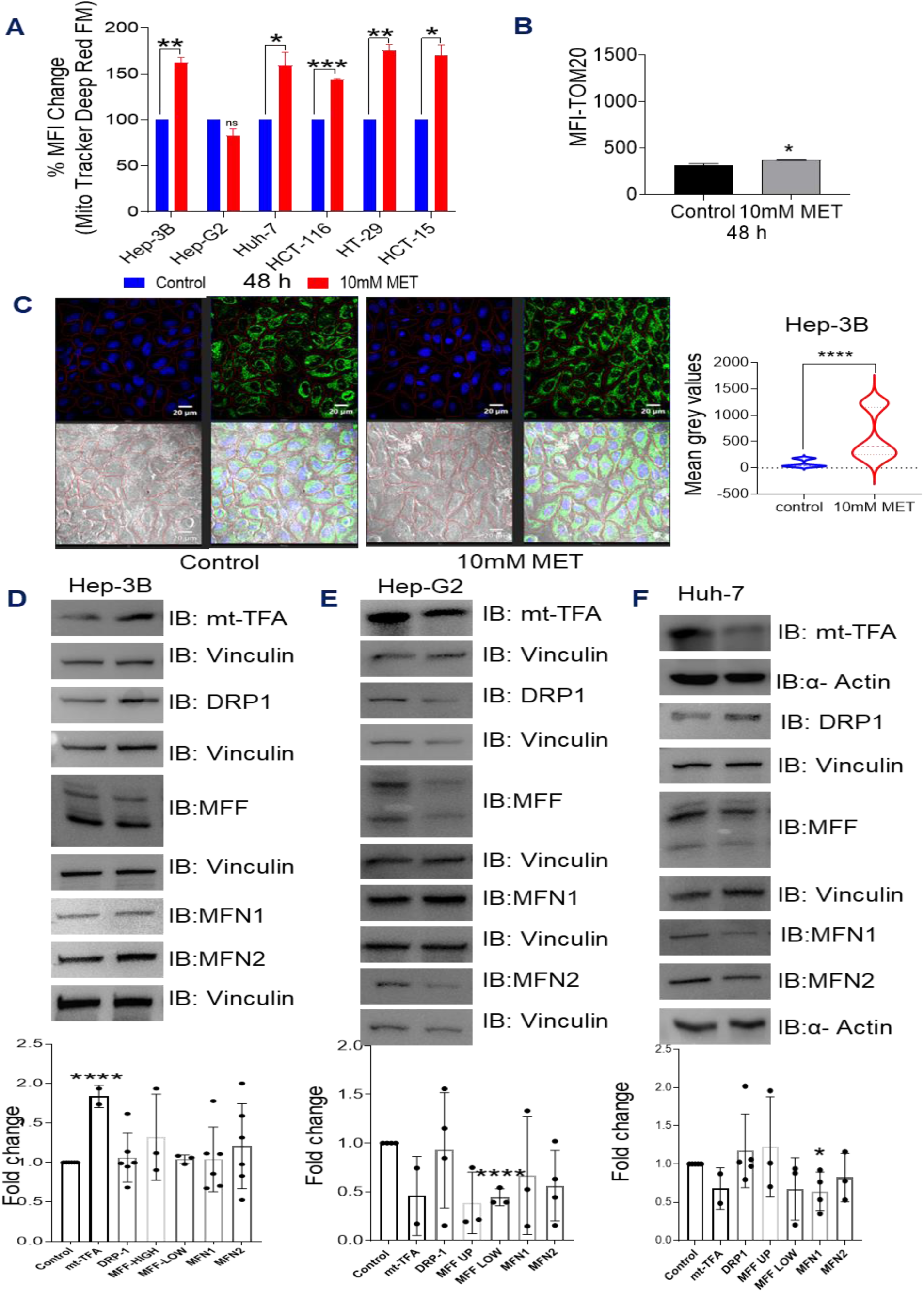
Effect of metformin on mitochondrial dynamics in human cancer cells. (A) HCC and CRC cells were treated with 10 mM metformin for 48 h and stained with MitoTracker Deep Red FM. Mean fluorescence intensity (MFI) was normalized to control (set as 100%), and results are shown as bar graphs representing mean ± S.D. from experimental replicates. (B) Hep-3B cells were treated with metformin for 48 h and stained for the mitochondrial marker TOM20. Flow cytometry data were acquired using FACS and analyzed with FlowJo software (N = 2). Bar graphs display mean ± S.D. from replicates of one representative experiment. (C) Hep-3B cells were treated with metformin for 36 h, stained for TOM20 (Green), and visualized by confocal microscopy. Single-cell MFI was quantified using CellSense software. Violin plots represent mean ± S.D. from replicates of one representative experiment (experiment repeated twice). (D–F) HCC cells (Hep-3B, Hep-G2, and Huh-7) were treated with metformin for 24 h. Expression of mitochondrial dynamics–related proteins were analyzed by western blot. Quantitative changes in protein expression were plotted as bar graphs representing mean ± S.D. Dots in the bar graphs indicate the number of independent experimental repeats.

In contrast, Hep-G2 cells reduced mt-TFA/TFAM together with decreased MFF and MFN1/2 was detected, while Huh-7 cells showed selective downregulation of mtTFA/TFAM and MFN1 with minimal induction of fission genes (Figure 1D-F). Analysis of publicly available RNA-seq datasets from metformin-treated HCC cells and PBMCs from healthy individuals administered metformin revealed differential effects on mitochondrial dynamics genes. In Huh-7 cells (GSE190076), 25 mM metformin for 48 h reduced transcript levels of fusion genes MFN1, MFN2, and OPA1, while fission-associated genes MFF, FIS1, and DNM1L were modestly increased. In Hep-G2 cells (GSE208245), metformin (0.25–3 mM) decreased fusion genes and selected fission components, along with downregulation of several mitochondrially encoded respiratory chain genes. Interestingly, In human PBMCs (GSE137317), non-significant changes were noted, suggesting that metformin exposure induces limited remodeling in normal circulating cells compared with HCC cells (Supplementary Figure S4 A-F).

These findings collectively indicate that metformin shifts mitochondrial dynamics toward increased fission in HCC cells, which contributes to metabolic reprogramming and adaptive remodeling in a cell line-dependent manner. Given the known roles of these molecules in regulating mitochondrial dynamics (Haq *et al*, 2013; Abu Shelbayeh *et al*, 2023), the results support that metformin treatment modulates key regulators of mitochondrial dynamics and metabolism. Moreover, the relative strength of fusion versus fission responses shapes mitochondrial plasticity and cancer cell fate under energetic stress.

### Metformin has minimal effect on tumor progression but alters mitochondrial markers level in tumors

Previous studies report that a metformin dose of 250 mg/kg/day in mice achieves a steady-state serum concentration of approximately 10 µM, sufficient to inhibit mitochondrial complex I. In humans, serum concentrations of ∼10–40 µM are attainable with 1–2 g/day dosing, with a maximum approved dose of 2.5 g/day due to good tolerability.

To evaluate metformin’s impact on tumor growth, two murine tumor models were used. The 4T1 triple-negative breast cancer model in immunocompetent BALB/c female mice, and the Hep-3B hepatocellular carcinoma model in immunocompromised NOD/SCID male mice were utilized. Tumor-bearing mice were randomized to receive metformin (250 mg/kg/day for 4T1, 250 and 500 mg/kg/day for Hep-3B) via drinking water or no treatment (Figure 2A). Tumor volume was measured every third day after palpable tumors appeared, with treatment continued for 18 days. Throughout the study, mice body weights remained stable, indicating minimal drug-related toxicity (Figure 2B and D). Surprisingly, metformin treatment did not significantly affect tumor growth in either model, as assessed by tumor volume measurements and excised tumor weights. The results indicate that metformin alone exerts minimal inhibitory effects on tumor progression in these preclinical models.

**Figure 2.**
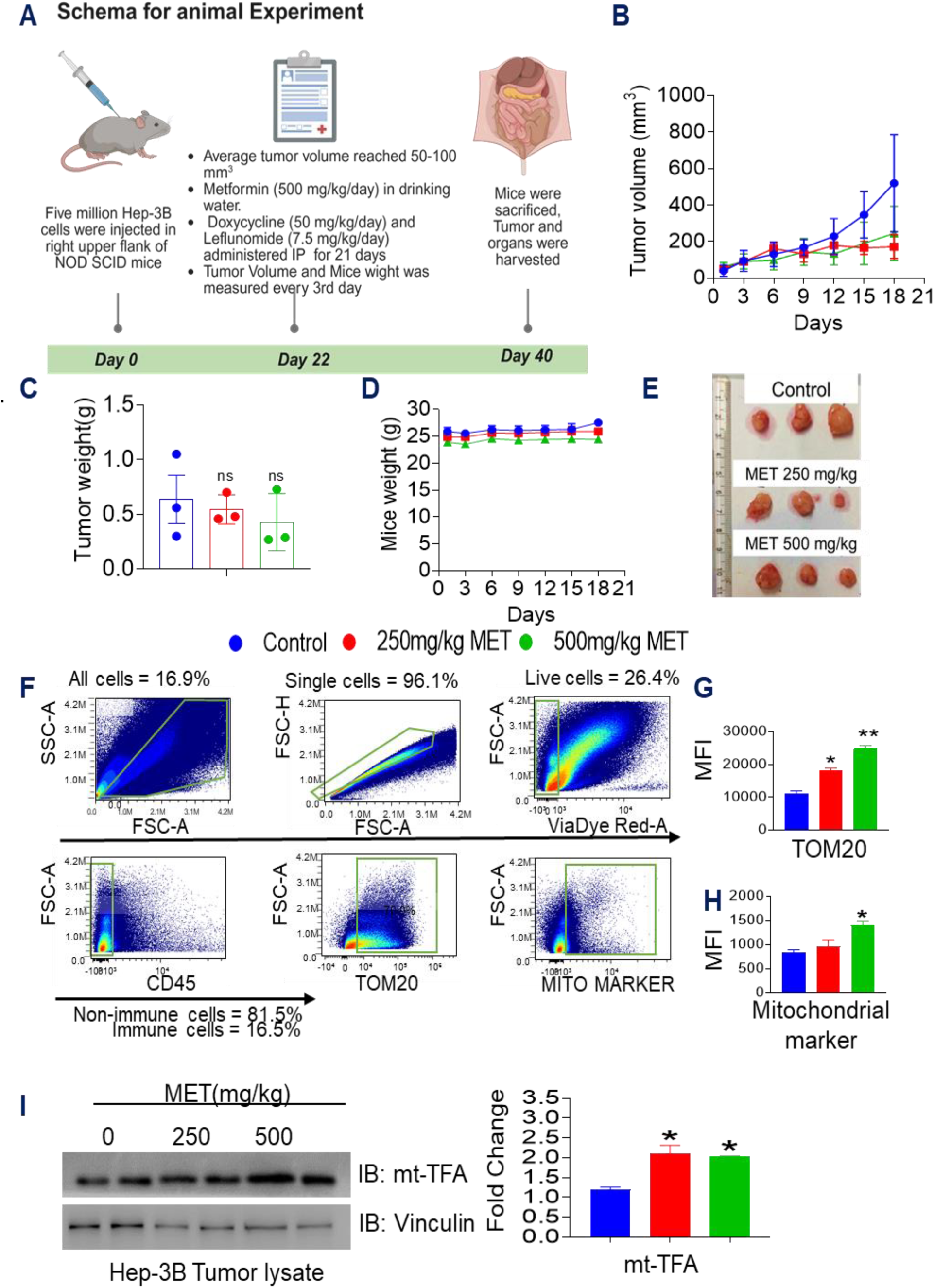
Effect of metformin on the growth of Hep-3B xenograft tumors and mitochondrial proteins. Hep-3B cells were injected subcutaneously into the upper right flank of NOD/SCID mice. NOD/SCID mice received metformin (250 or 500 mg/kg/day) in drinking water throughout the experimental duration. At the end of treatment, tumors were excised, weighed, and processed to prepare single-cell suspensions. (A) Schematic representation of the animal study design (created with BioRender, https://BioRender.com/t62p562). (B) Tumor growth curve showing changes in Hep-3B xenograft volume over time. (C) Tumor weights at the experimental endpoint. (D) Body weight of tumor-bearing mice recorded throughout the study. (E) Representative photographs of excised Hep-3B tumors. These tumor samples were used for single-cell suspension preparation for flow cytometry and western blot analysis. (F) Flow cytometric gating strategy used to evaluate mitochondrial markers in single-cell suspensions. (G) Mean fluorescence intensity (MFI) of TOM20. (H) MFI of additional mitochondrial markers analyzed. (I) Representative western blot and quantitative analysis of mean gray values for mt-TFA expression. Lysates were prepared from single-cell suspensions, and images were captured using a UVTECH ChemiDoc and analyzed with ImageJ software. All quantitative results were plotted using GraphPad Prism.

To determine whether metformin nonetheless modulates tumor mitochondrial biology, excised Hep-3B tumors were processed into single-cell suspensions for flow cytometry and western blot analysis. Representative photographs of tumors used for these assays are shown in Figure 2E. Using the gating strategy outlined in Figure 2F to focus on live, non-immune (CD45⁻) tumor cells, metformin treatment was associated with increased TOM20 mean fluorescence intensity (MFI), indicating enrichment of mitochondrial outer membrane protein relative to controls (Figure 2G). Status of pan-mitochondrial marker showed a similar increase in MFI, supporting an overall rise in mitochondrial signal and suggesting augmented mitochondrial content or mass within metformin-treated tumors (Figure 2H). Consistent with this, immunoblotting of tumor lysates revealed a dosedependent elevation of mitochondrial transcription factor A (mt-TFA/TFAM), a key regulator of mtDNA maintenance and mitochondrial biogenesis, in metformin-treated groups (Figure 2I). A similar lack of effect on primary tumor growth was observed in the independent 4T1 isograft model, where tumor volumes, endpoint weights, and body weights were indistinguishable from controls (Supplementary Figure S5 A–E). Analysis of 4T1 tumors from metformin-treated mice revealed a modest but consistent increase in TOM20 MFI compared with controls (Supplementary Figure S5 F and G). Taken together, these findings indicate that while metformin does not significantly constrain tumor growth in the models tested, it consistently promotes mitochondrial remodeling in vivo, characterized by enhanced mitochondrial marker expression and upregulation of mt-TFA, consistent with an adaptive increase in mitochondrial content and transcriptional capacity within tumor cells.

### Targeting of metformin-promoted changes in mitochondrial dynamics induces cytotoxicity in cancer cells

Above mentioned results indicate that metformin treatment modulates mitochondrial biogenesis and fission. To assess the significance of these processes for cancer cell survival, we used pharmacological agents targeting mitochondrial biogenesis and fission (Scatena *et al*, 2018). Doxycycline, which inhibits mitochondrial biogenesis, and Mdivi-1, a selective DRP1 inhibitor that blocks mitochondrial fission, were used. Leflunomide, a DHODH inhibitor that promotes mitochondrial fusion was also evaluated (Alamri *et al*, 2021).

Analysis of fusion–fission related transcripts after 24 h indicated that Mdivi-1 treatment alone upregulated MFF1 and FIS1, with modest increases in DRP1, MFN1 and MFN2 mRNA in Hep-3B cells, while inducing only minor, non-significant changes in Huh-7 cells. Leflunomide modestly increased MFN1 and MFN2 expression in Hep-3B and caused only slight changes in Huh-7, suggesting DHODH inhibition exerts a relatively mild pro-fusion signal in these HCC models (Supplementary Figure S3A-D).

In cell death assays, metformin, doxycycline, Mdivi-1, or leflunomide alone caused only modest increases in cell death at 36–48 h. In contrast, combining each mitochondrial modulator with metformin produced pronounced cytotoxicity, with metformin + doxycycline, metformin + Mdivi-1 and metformin + Leflunomide consistently yielding the highest levels of cell death in both Hep-3B and Huh-7 cells (Figure 4A–D). These data demonstrate strong relationship between complex I inhibition and interference with mitochondrial translation, biogenesis, or the fission/fusion machinery in cells that have undergone metformin-induced mitochondrial remodeling.

Mechanistically, combined metformin + Mdivi-1 treatment in Hep-3B cells strongly impaired bioenergetic flexibility. Basal oxygen consumption rate, proton leak, metabolic potential, spare respiratory capacity, and lactate production per glucose were all reduced, shifting cells into a quiescent, low-flexibility energetic state (Supplementary Figure S6A– G). Notably, co-treatment also robustly activated lysosomal function, as evident by increased acidified lysosomal compartments and higher LysoSensor fluorescence. Disruption of lysosomal function using bafilomycin or leupeptin attenuated the cytotoxic and growth-inhibitory effects, indicating that autophagy–lysosomal flux plays a key role in executing cell death under dual mitochondrial stress (Supplementary Figure S7A-D). Consistent with this, co-treatment with the antioxidant N-acetylcysteine partially rescued growth inhibition in Mdivi-1- or metformin + Mdivi-1-treated Hep-3B and Huh-7 cells, implicating a role of reactive oxygen species in promoting cytotoxicity (Figure 4E-F). Overall, these data demonstrate that targeting mitochondrial adaptations induced by metformin, particularly biogenesis and fission processes, enhances cytotoxicity in liver cancer cells by exploiting mitochondria-dependent vulnerabilities that can be targeted by perturbing mitochondrial translation, fission, DHODH-linked fusion pathways, and lysosomal function.

### Inhibiting metformin-induced changes in mitochondrial dynamics restricts tumor growth

Our findings indicate that metformin alone exerts minimal inhibitory effects on solid tumor growth. Based on in vitro results demonstrating that metformin combined with doxycycline or leflunomide induces significant death in HCC cells, we evaluated the therapeutic efficacy of these combinations in vivo.

Hep-3B tumor-bearing mice were randomized into control and five experimental groups (4 to 6 animals each): Group 1, metformin; Group 2, leflunomide; Group 3, metformin plus leflunomide; Group 4, doxycycline; and Group 5, metformin plus doxycycline (Figure 4A1). Metformin was delivered in drinking water, while leflunomide and doxycycline were administered via intraperitoneal injection. Control mice received vehicle solution. Tumor progression was monitored by measuring tumor volume every third day. No significant weight changes were observed in any treatment groups throughout the study, indicating low systemic toxicity (Figure 4C). Liver and kidney histology studies of control and metformin + leflunomide animals showed no abnormalities, while metformin or leflunomide alone caused only minimal hepatic changes. Doxycycline treatment caused mild hepatocellular alterations, which were reduced to minimal in the metformin + doxycycline group. Renal sections in all groups were graded as normal or with only minimal focal tubular degeneration.

Single-agent treatments with metformin, doxycycline, or leflunomide did not significantly reduce tumor volume compared to controls, with tumors retaining a moderately developed, highly cellular morphology and only minimal focal necrosis. In contrast, combination treatments with metformin plus leflunomide or metformin plus doxycycline significantly suppressed tumor growth compared to single agents (Figure 5B and 5D). Histopathological grading of tumor sections revealed that leflunomide alone induced minimal (+1) degenerative and necrotic changes, whereas metformin + leflunomide increased these to mild (+2) with multiple necrotic foci, and metformin + doxycycline produced the most pronounced effect with moderate (+3) widespread necrosis and destruction of tumor tissue. (Supplementary Figure S8 A-F, Supplementary Table 1).

These in vivo findings, together with the in vitro data, suggest that co-targeting mitochondrial translation or DHODH in metformin-primed cells disrupts metabolic flexibility and triggers cell death. Collectively, these data demonstrate that targeting metformin-induced mitochondrial adaptations with inhibitors of biogenesis or fission effectively restricts tumor growth in vivo while causing only minimal, largely reversible changes in major organs, highlighting the potential of combined therapeutic strategies to potentiate metformin’s anticancer efficacy.

## Discussion

Metformin’s role as an anticancer agent has been extensively investigated, yet the accumulated evidence from preclinical and clinical studies remains equivocal. A rigorous meta-analysis encompassing 22 randomized controlled trials with 5,943 cancer patients revealed that metformin monotherapy or combined with standard treatments yielded predominantly neutral outcomes (∼64% trials) and only modest positive effects (∼23%) largely confined to reproductive system cancers. Conversely, in digestive system malignancies such as pancreatic and liver cancers, metformin may even worsen progression-free survival in some cases (Wen *et al*, 2022; Zhang *et al*, 2022). This inconsistency in efficacy aligns with our findings showing minimal tumor growth inhibition in 4T1 breast cancer and Hep-3B liver cancer xenografts with metformin alone, despite achieving plasma concentrations commensurate with mitochondrial complex I inhibition (Figure 2). Such observations are in line with reports that metformin concentrations tolerable in humans are insufficient to induce substantial cancer cell death in vitro (He & Wondisford, 2015; Mogavero *et al*, 2017).

Our data indicate that metabolic plasticity and mitochondrial adaptive remodeling in cancer cells markedly constrain metformin’s antitumor efficacy. In Hep-3B cells, metformin increased mitochondrial biomass and altered dynamics while maintaining residual respiration and glycolytic compensation, similar to OXPHOS-adapted trajectories described in other metformin-treated tumors. These adaptive mitochondrial dynamics, reflected in increased expression of TFAM, MFF, and TOM20 in vitro and in vivo (Figures 1, 2), appear to help cancer cells withstand metformin-induced metabolic insults, thereby maintaining survival and tumor growth.

Disrupting this metformin-primed state exposes targetable mitochondrial vulnerabilities. Combining metformin with inhibitors of fission (Mdivi-1), DHODH-dependent metabolism (leflunomide), or mitochondrial translation (doxycycline) markedly increased cell death and reduced viability compared with single agents in Hep-3B and Huh-7 cells (Figure-3). These combinations abrogate basal and spare respiratory capacity with limited glycolytic rescue, increase ROS levels, and activate lysosome-dependent pathways leading to enhanced cytotoxic effect (Supplementary Figure- S6)(Figure-3G-H). Interestingly, cytotoxicity induced by combinations was partially reversed by lysosomal or ROS scavenging, indicating that metformin-remodeled mitochondria become highly sensitive to combined energetic, oxidative, and proteotoxic stress (Supplementary Figure S7). Additionally, inhibition of DRP1 or DHODH is also likely to impair mitophagy, which can lead to accumulation of dysfunctional, metformin-damaged mitochondria, thereby further amplifying stress.

Targeting mitochondrial dynamics emerged as a promising strategy to sensitize cancer cells to metformin. Pharmacologic inhibition of biogenesis (doxycycline), fission (Mdivi-1), or promotion of fusion (leflunomide) synergistically enhanced metformin-induced cytotoxicity in HCC cells (Figure 3). Bioenergetic analyses revealed that combinations impair metabolic flexibility more effectively than single agents, as evidenced by decreased basal respiration and extracellular acidification rates (Supplementary Figure S6). These findings underscore mitochondrial dynamics as critical nodes of metabolic vulnerability that, when co-targeted, potentiate metformin’s antitumor effects in vitro and in vivo (Figure 3 and 4).

**Figure 3.**
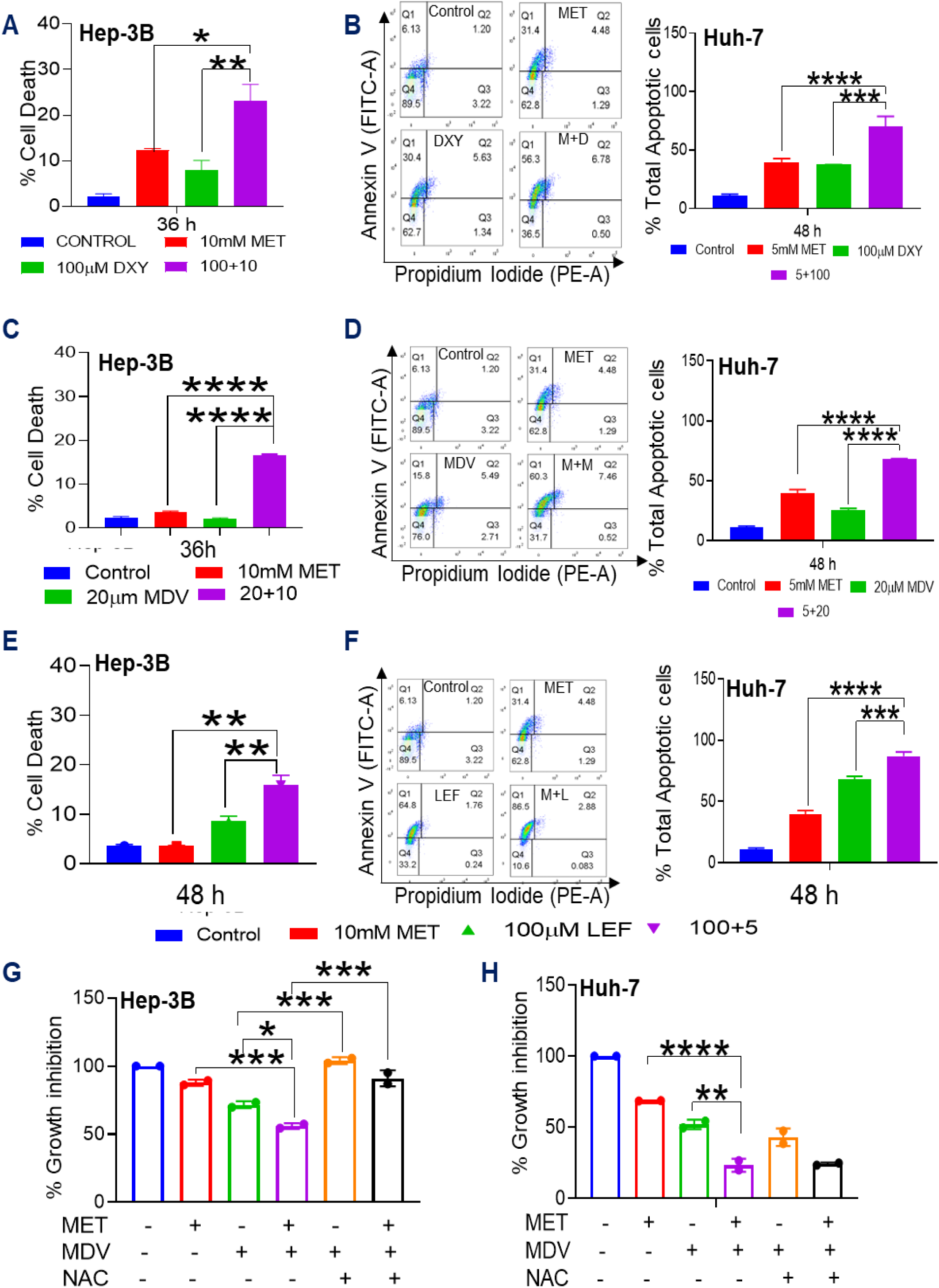
Targeting metformin-induced mitochondrial remodeling in cancer cells. Equal numbers of Hep-3B, Huh-7, and Hep-G2 cells were seeded and treated with metformin alone or in combination with mitochondrial dynamics modulators. Cell death was assessed in Hep-3B cells by propidium iodide (PI) exclusion and in Huh-7 cells by Annexin V/PI staining, followed by flow cytometry. (A) Hep-3B and (B) Huh-7 cells treated with metformin and doxycycline (DXY), alone or in combination, for 36 h or 48 h, respectively. (C) Hep-3B and (D) Huh-7 cells treated with metformin and Mdivi-1 (MDV), alone or in combination, for 36 h or 48 h, respectively. (E) Hep-3B and (F) Huh-7 cells treated with metformin and leflunomide (LEF), alone or in combination, for 48 h. PI-positive or Annexin V/PI-stained populations were quantified using FlowJo. (G) Hep-3B and (H) Huh-7 cells treated for 48 h with metformin (10 mM) alone or in combination with MDV (20 µM), MDV plus N-acetylcysteine (NAC), or metformin plus MDV plus NAC, followed by crystal violet staining and percentage growth inhibition was quantified using Fiji (ImageJ).

**Figure 4:**
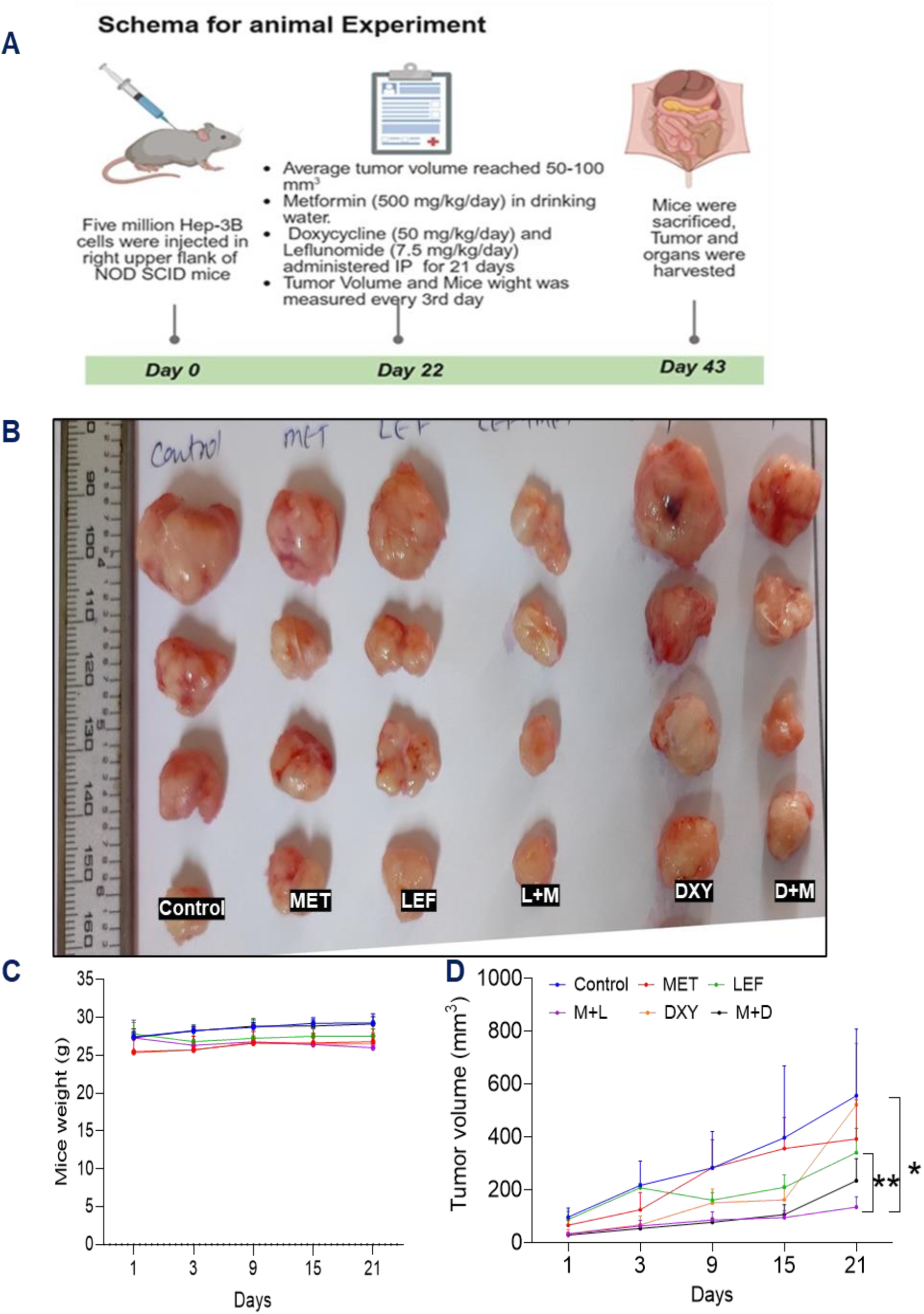
Targeting metformin-induced mitochondrial dynamics inhibits Hep-3B xenograft tumor growth. Hep-3B cells were injected into NOD/SCID mice, which were treated with metformin (500 mg/kg/day), doxycycline (50 mg/kg/day), and leflunomide (7.5 mg/kg/day) individually or in combination as described in the Methods. At endpoint, tumors were excised and analyzed. (A) Experimental schematic of Hep-3B xenografts in NOD/SCID mice (illustration created with BioRender, https://biorender.com/t62p562). (B) Representative photographs of excised tumors. (C) Body weights of tumor-bearing mice during treatment. (D) Tumor volumes measured longitudinally using digital vernier calipers.

In vivo, metformin combined with leflunomide or doxycycline significantly reduced Hep3B xenograft tumor size relative to control and monotherapies, with extensive tumor necrosis but preserved body weight and only minimal, largely reversible hepatic and renal changes, supporting a favorable short-term therapeutic window (Figure-4) (Supplementary Figure S8) (Supplementary Table-1). These data, together with the in vitro results, suggest that metformin functions as a mitochondrial priming agent. It causes expansion and reconfiguration of the mitochondrial network which creates a state that remains viable alone but becomes synthetically lethal when fission, DHODH-linked respiration, or mitochondrial translation are additionally blocked.

While these results are compelling, limitations should be acknowledged. This study has limitations including reliance on a small panel of liver cancer cell lines, context-dependent resistance in less adaptable models such as Hep-G2, use of immunodeficient xenografts that omit immune and stromal effects, and single-time-point analyses that may miss dynamic regulation of mitochondrial proteins and transcripts. Moreover, well-designed clinical trials integrating metabolic modulators alongside metformin, supported by mechanistic biomarker studies, are essential to harness its full anticancer potential. In summary, our study reveals that metformin-induced mitochondrial remodeling limits its anticancer efficacy by fostering metabolic adaptation and survival, but this creates exploitable vulnerabilities. Co-targeting mitochondrial biogenesis and fission disrupts this adaptive plasticity, inducing metabolic collapse and tumor regression. This mechanistic insight provides a rationale for combinational therapeutic approaches aimed at overcoming metabolic resistance and enhancing metformin’s clinical utility in cancer therapeutics.

## Work Summary: Cancer cells alter metabolism to hinder metformin’s ability to decrease cell survival

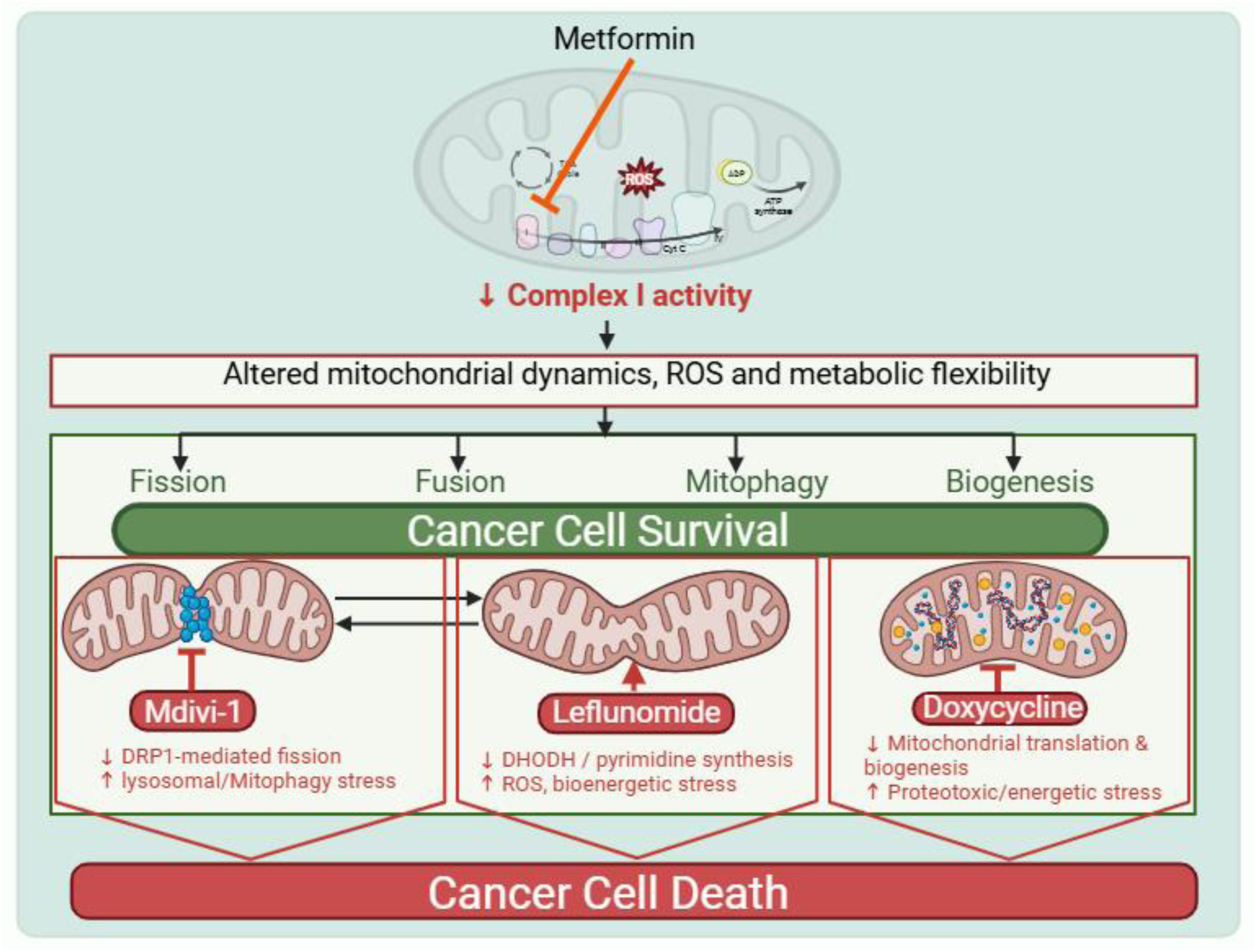

## Ethics approval

All the animal experiments were performed as per the requirement and guidelines of the Committee for the Purpose of Control and Supervision of Experiments on Animals (CPCSEA), Government of India, and after obtaining permission from the Institutional Animal Ethics Committee (IAEC; IAEC/2018/B-335, IAEC/2024/B-473).

## Declaration of Generative AI and AI-assisted technologies in the writing process

The authors used an AI-based language model (Perplexity AI) to assist in improving the grammar and readability of the manuscript. The final version was thoroughly revised and approved by all authors to ensure scientific accuracy and integrity.

## Acknowledgements

The author thank former directors of NCCS Dr. S.C. Mande, and late Dr. Mohan R. Wani, NCCS Pune India for being very supportive and giving all the encouragement to carry out this work. F.K.B. and U.S. acknowledges University Grants Commission (UGC), New Delhi, India for research fellowship. F.K.B also acknowledges the Science and Engineering Research Board, Department of science and technology for senior research fellowship. The authors thank Biotechnology Research and Innovation Council - National Centre for Cell Science (BRIC- NCCS), Pune, India, and Savitribai Phule Pune University, Pune, India for their support.

## Author contributions

F.K.B: Conceptualization, Formal Analysis, Investigation, Methodology, Validation, Visualization, Writing – original draft, Writing – review & editing. M.K.B.: Conceptualization, Funding acquisition, Methodology, Project administration, Resources, Supervision, Writing – review & editing. U.S.: Writing – review & editing, Methodology, Formal Analysis. S.S: Resources, Methodology, Formal Analysis, Writing – review & editing.

