## Supplementary Material for "Metformin-Induced Mitochondrial Remodeling Creates Adaptive and Targetable Vulnerabilities in Cancer Cells"

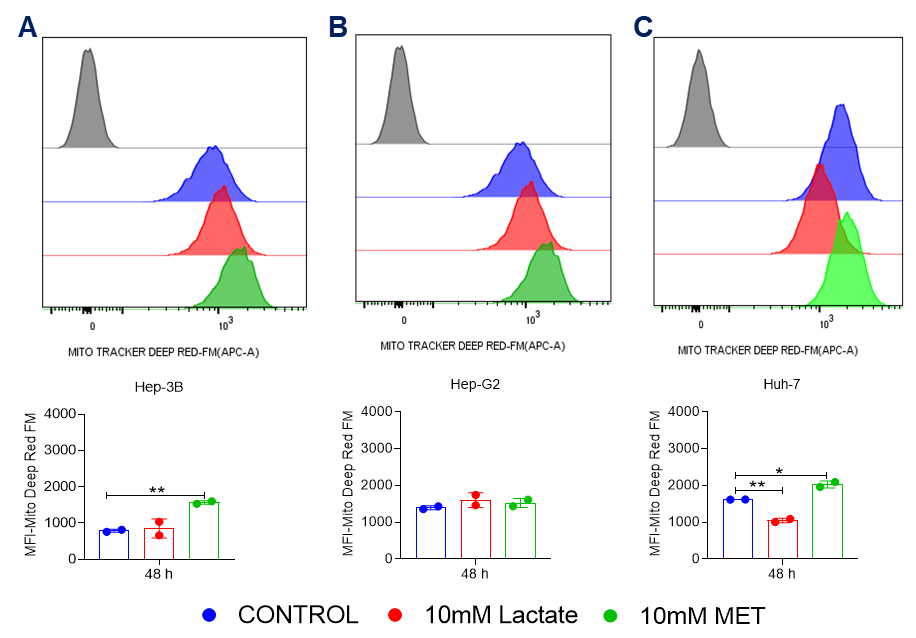

**Supplementary Figure S1. Effect of metformin and lactate on mitochondrial density in Hep‑3B, Hep‑G2, and Huh‑7 cells.**
Equal numbers of cells were seeded and allowed to attach for 24 h, then treated with metformin (10 mM) or lactate (10 mM) for 48 h. At the end of treatment, cells were labeled with MitoTracker Deep Red FM and analyzed by flow cytometry. Histograms and mean fluorescence intensity (MFI) values are shown for (A) Hep‑3B, (B) Hep‑G2, and (C) Huh‑7 cells. Data were processed using FlowJo, and graphs and statistical analyses were generated in GraphPad Prism using unpaired Student’s t‑test; p‑values are indicated as *p < 0.05, **p < 0.01, ***p < 0.001, ****p < 0.0001.

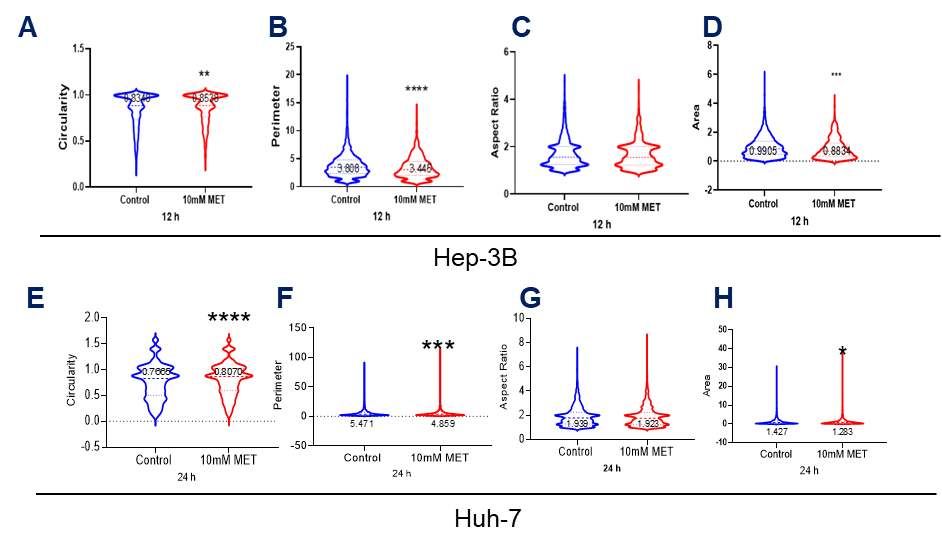

**Supplementary Figure S2. Effect of metformin on mitochondrial morphology in Hep‑3B and Huh‑7 cells.**
Equal numbers of cells were seeded on coverslips in 24‑well plates and allowed to attach for 24 h, then treated with metformin for the indicated time points. At the end of treatment, Hep‑3B cells were stained for TOM20 and Huh‑7 cells were labeled with MitoTracker Deep Red FM. Confocal images were acquired and mitochondrial morphology was quantified in ImageJ (Fiji) using mitochondrial analyzer plugins. (A, E) Circularity, (B, F) perimeter, (C, G) aspect ratio, and (D, H) area of mitochondrial structures are shown. Graphs and statistical analyses were generated in GraphPad Prism using unpaired Student’s t‑test; p‑values are indicated as *p < 0.05, **p < 0.01, ***p < 0.001, ****p < 0.0001.

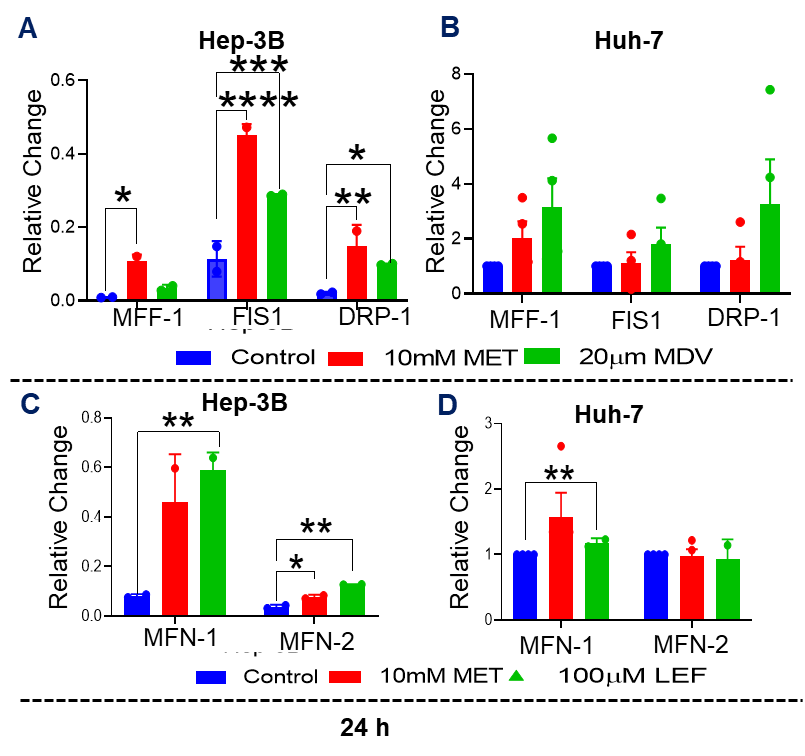

**Supplementary Figure S3. Effect of mitochondrial dynamics modulators on expression of mitochondrial fission- and fusion-related genes in Hep‑3B and Huh‑7 cells.**

Equal numbers of cells were seeded, allowed to attach for 24 h, and then treated with metformin (MET, 10 mM), Mdivi‑1 (MDV, 20 µM), or leflunomide (LEF, 100 µM) for an additional 24 h. At the end of treatment, mRNA expression of fission-related genes MFF1, FIS1, and DRP1 was quantified in (A) Hep‑3B and (B) Huh‑7 cells, and fusion-related genes MFN1 and MFN2 were measured in (C) Hep‑3B and (D) Huh‑7 cells. Graphs and statistical analyses were generated in GraphPad Prism using unpaired Student’s t‑test; p‑values are indicated as *p < 0.05, **p < 0.01, ***p < 0.001, ****p < 0.0001.

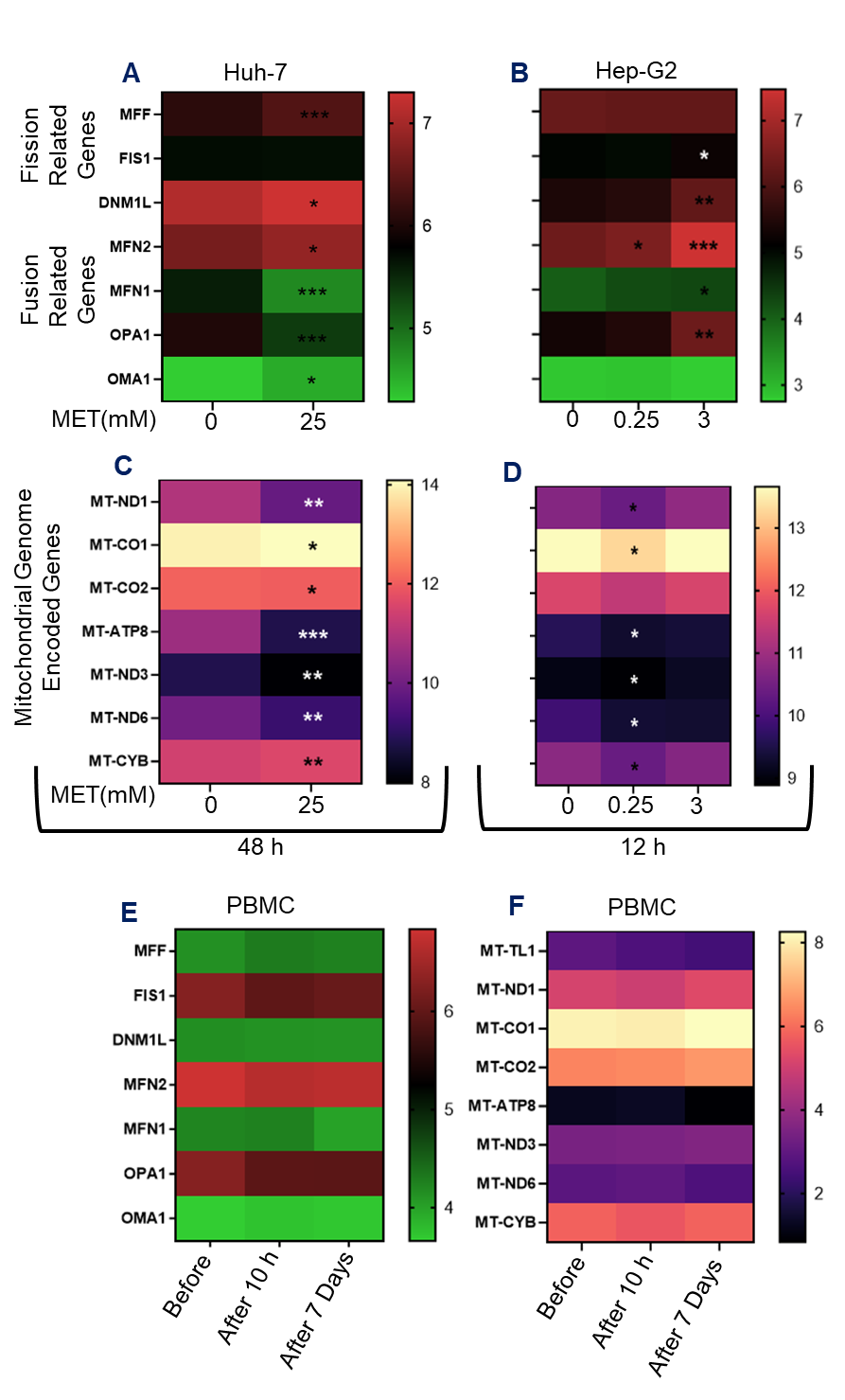

**Supplementary Figure S4:** **Effect of metformin treatment on the gene expression in HCC cell line and normal human PBMC**. Data were acquired from GEO (Huh-7; GSE190076, Hep-G2; GSE208245, PBMC; GSE137317) and analyzed as described in methodology section. Human HCC cells and PBMC were treated with metformin and RNAseq was done. The available data were analyzed for the change in the expression of mitochondrial dynamics related genes and mitochondrial genome encoded genes expression. Color gradient graphs showing changes in mitochondrial dynamics related genes in (A) metformin (25 mM) treated Huh-7 cells compared to untreated. (B) Metformin treated Hep-G2 cells (0.25 and 3 mM). Color graph showing changes in mitochondrial genome encoded gene expression in (C) Huh-7 cells, (D) Hep-G2 cells. Normal human individuals were treated with metformin and their PBMC were analyzed for the changes in the expression of (E) mitochondrial dynamics related genes and (F) mitochondrial genome encoded gene expression. *Graphs and statistical analysis were done using GraphPad Prism. For statistical analysis, a paired t-test was performed, and p-values are denoted as follows: *p < 0.05, **p < 0.01, ***p < 0.001, and ****p < 0.0001.*

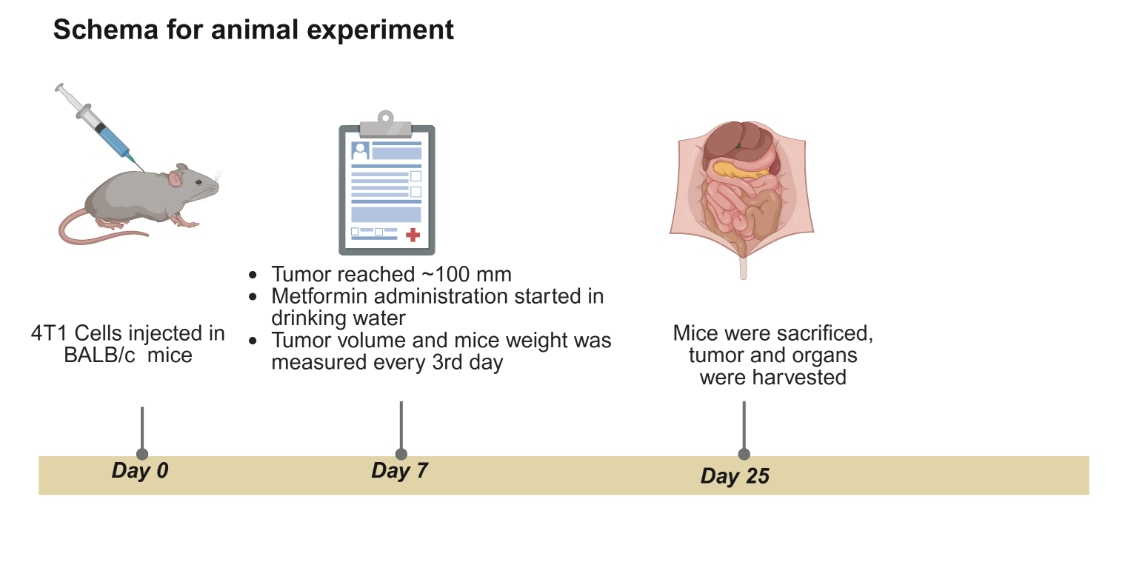

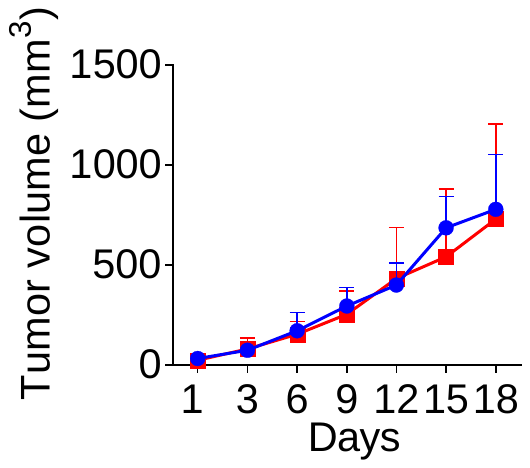

**A**

**B**

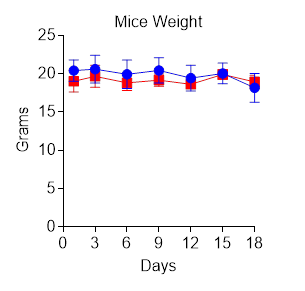

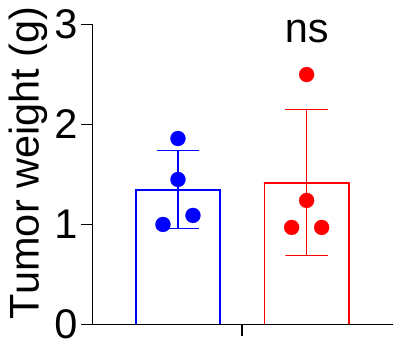

Control

**E**

**D**

**C**

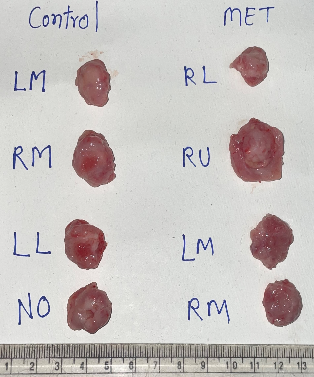

Metformin

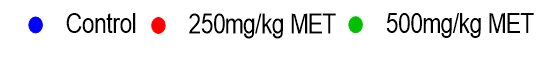

**F**

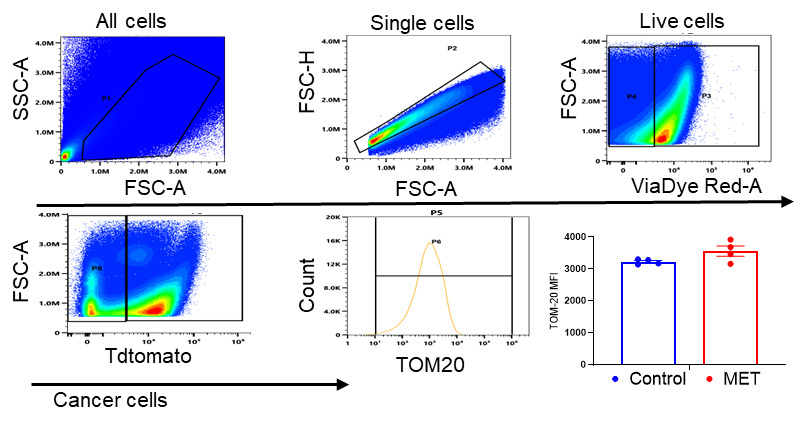

**G**

**Supplementary Figure S5: *Effect of metformin treatment on mitochondrial biomass in 4T1 isograft tumor cells.*** *4T1 cells were injected into mammary fat pad of Balb/c mice, and treatments of 250 mg/kg metformin was provided through drinking water. (A) Schema of animal experiment (Created in BioRender. https://BioRender.com/t62p562), (B) 4T1 tumor volume measured by digital vernier caliper, (C) 4T1 tumor photograph (D) Excised 4T1 tumor weight, (E) Mice weight during the treatment. At the end of the experiment, tumors were excised, and single-cell suspensions were prepared. Cells were stained with respective markers an analyzed by using FACS (F) Shows gating strategies for the evaluation of mitochondrial marker TOM20 in single-cell suspension cells in which single cells were gated for live and dead cells by using ViaDye Red, ViaDye Red negative cells were gated for the Tdtomato (4T1 cells expresses this protein) which separate cancer cells from other cells. (G) The MFI of TOM-20 in 4T1 tumor cells were calculated by analyzing data using SpectroFlo and OMIQ software. Graphs and statistical analysis were done using GraphPad Prism. For statistical analysis, an unpaired Student’s t-test was performed, and p-values are denoted as follows: *p < 0.05, **p < 0.01, ***p < 0.001, and ****p < 0.0001.*

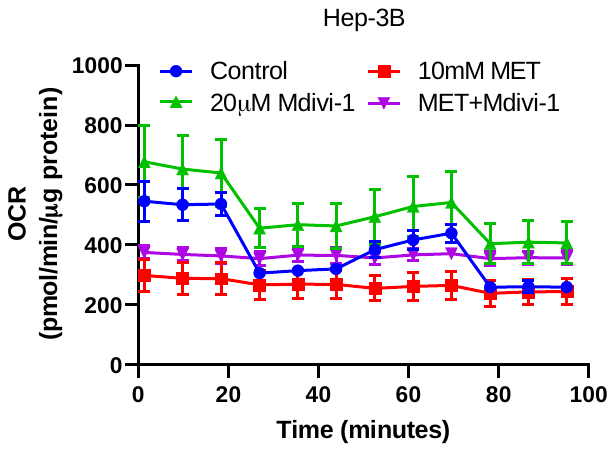

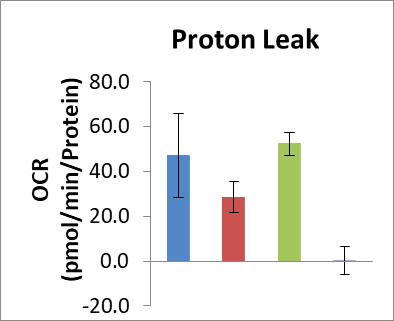

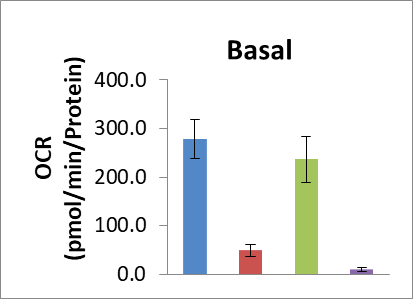
**A	B	C**

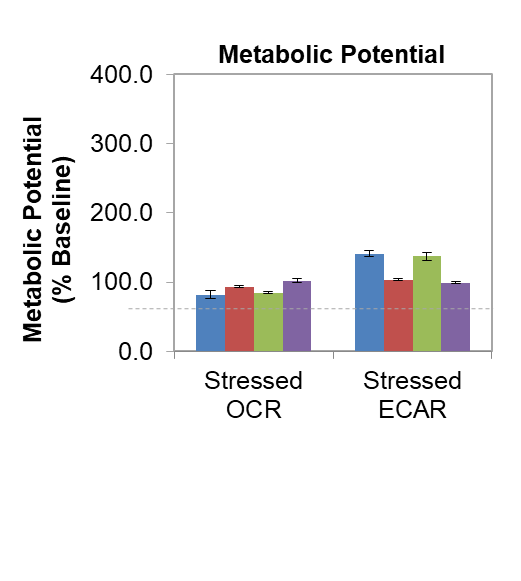

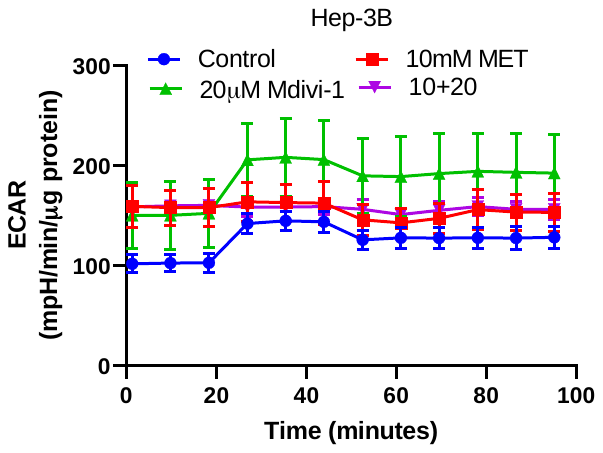

**D	E**

**G**

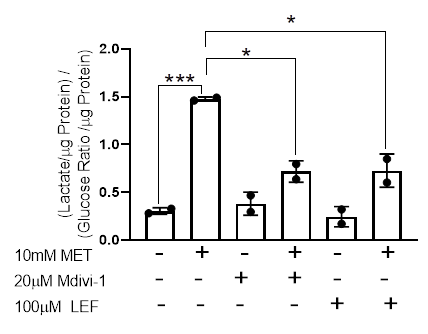
 **F**

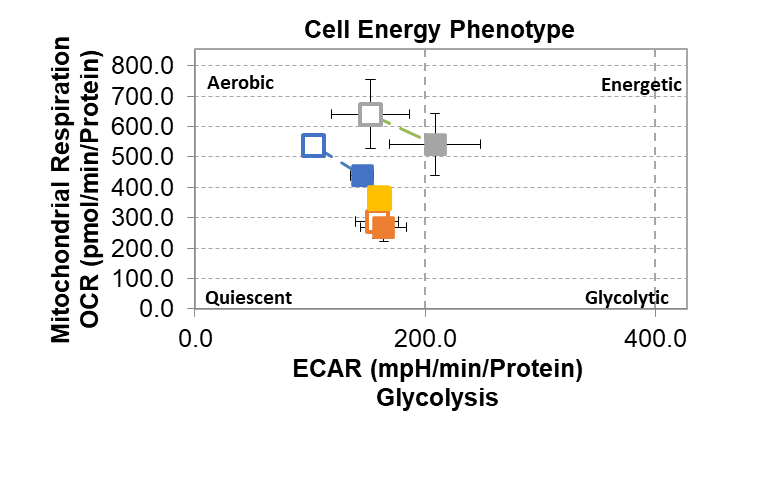

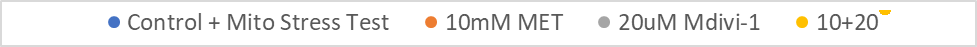

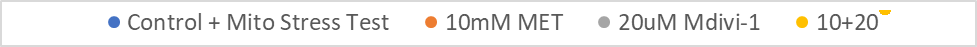

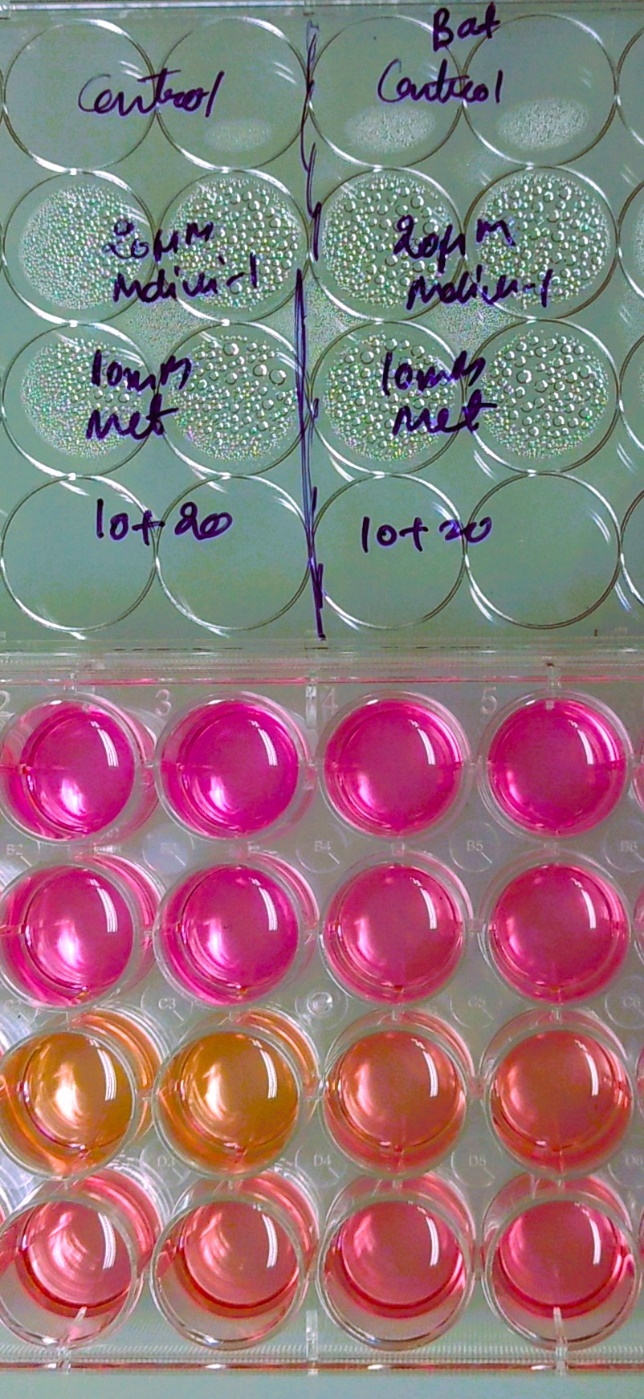
**Supplementary Figure S6:** **Effect of combine targeting of mitochondrial complex-1 and fission on cellular energetics of Hep-3B cells.** Hep-3B cells were treated with inhibitors alone or in combination for 24 h and subjected to the bioenergetic phenotype testing afterward the data were normalized with protein concentrations in each well graphs are showing (A) Oxygen consumption rate (OCR) (B) Basal respiration rate (C) Proton leak, (D) Extracellular acidification rate (ECAR), (E) Metabolic Potential and (F) cell energy phenotype. (G) Change in the ratio of glucose and lactate in spent media.

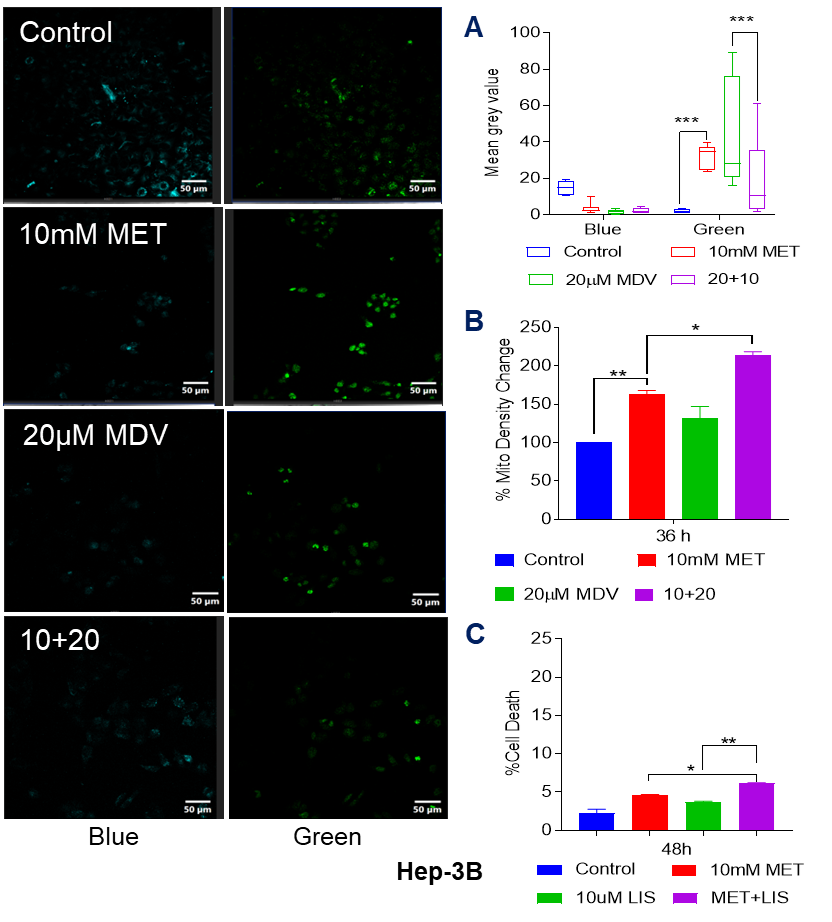

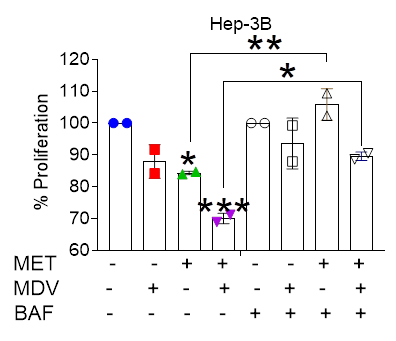

**D**

**Supplementary Figure S7: Effect of complex I and DRP-1 inhibition on the activation of lysosomes.** Cells were seeded equally on glass coverslips, and after 24h, treatments of respective drugs were administered. After treatment, cells were stained with LysoSensor Yellow/Blue DND-160 and acquired using an Olympus 3000 confocal microscope as per the protocol mentioned in the methodology section. Images are displayed with added coloring showing blue/cyan for neutral lysosomes and green for activated lysosomes in (A). To quantify the expression, each cell was individually selected as a region of interest, and mean grey values were calculated using ImageJ Software (B). Similar treatments were administered to evaluate Mitotracker Deep Red FM median fluorescence intensity, which was calculated (C). Treatment of MET and liensinine were given to Hep-3B cells, and PI exclusion cell death assay was performed. (D) Hep‑3B cells were treated for 36 h with metformin (MET, 10 mM), Mdivi‑1 (MDV, 20 µM), and/or bafilomycin A1 (BAF, 10 nM) as indicated. Cell proliferation/viability was quantified at the end of treatment by MTT assay and expressed as percent of vehicle control. Data was analyzed by FlowJo Software after FACS acquisition. For statistical analysis, two-way ANOVA with Tukey’s multiple comparisons were performed for lysosome staining, and ordinary one-way ANOVA was performed for mitochondrial staining. P-values are denoted by “,” where a p-value <0.05(), <0.01(), <0.001() and <0.0001(***).

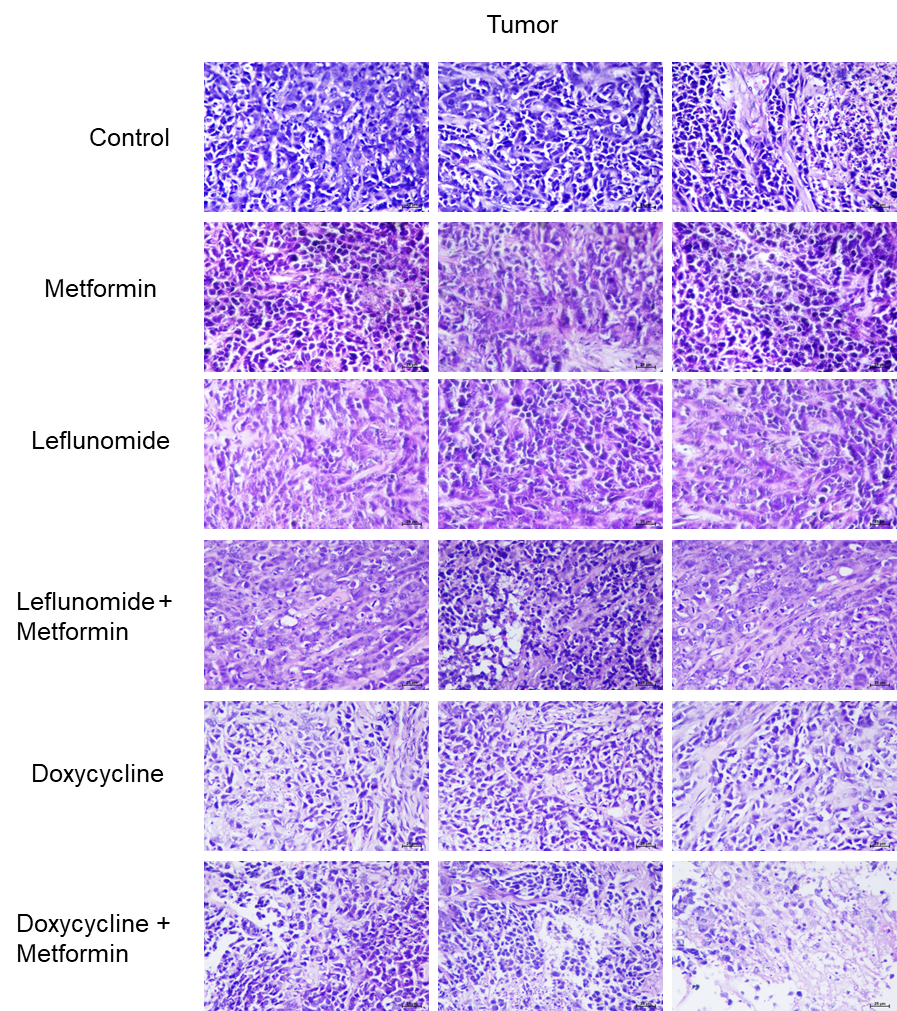

**Supplementary Figure S8:** **Effect of metformin, leflunomide, doxycycline on Hep-3B tumor histopathology either as single drug or in combination with metformin**. H & E staining was done and images were taken. Images were analyzed by histopathologist and interpretation given are mentioned in supplementary Table 1.

**Supplementary Table-1: Histopathological observations on sections of organs and tumor harvested from mice treated with MET (Metformin), LEF(Leflunomide), DXY(Doxycycline) as single drug or in combination with respective co**

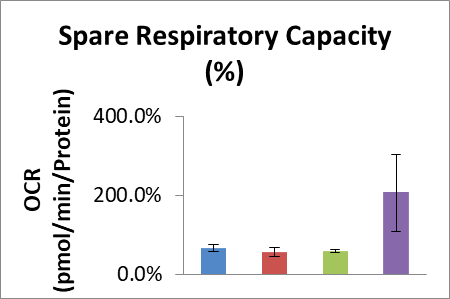

| **Sr. N.** | **Group /Slide Code** | A. **Histopathological observations**   **Tumour Tissue** | **Grade for tumour cell destruction** |
| --- | --- | --- | --- |
| **1** | **Control** | Moderately developed tumor tissue of proliferative fibrocytes and fibroblasts.  The tumour tissue comprised of variable sized cells of elongated and spindle shape with dense arrangement.  Prominent basophilia with enlarged nuclei, mitotic figures and marked pleomorphism of fibroblastic neoplastic cells was noted. | **Non-significant & focal** |
| **2** | **MET** | Moderately developed tumor tissue of proliferative fibrocytes and fibroblasts. The tumour tissue comprised of variable sized cells of elongated and spindle shape with dense arrangement.  Prominent basophilia with enlarged nuclei, mitotic figures and marked pleomorphism of fibroblastic neoplastic cells was noted. | **Non-significant & focal** |
| **3** | **LEF** | The tumour tissue comprised of variable sized cells of glands and fibroblasts as major cellular component. The tumour tissue comprised of variable sized cells of elongated and spindle shape with dense arrangement. Prominent basophilia with enlarged nuclei, mitotic figures and marked pleomorphism of fibroblastic neoplastic cells was noted.  **Few foci showed degenerative and necrotic changes of neoplastic cells in the tumour mass resulting into cellular debris.** | **(+1)**  **Minimal** |
| **4** | **MET - LEF** | The tumour tissue comprised of variable sized cells of glands and fibroblasts as major cellular component. The tumour tissue comprised of variable sized cells of elongated and spindle shape with dense arrangement. Prominent basophilia with enlarged nuclei, mitotic figures and marked pleomorphism of fibroblastic neoplastic cells was noted.  **Multiple foci showed marked necrotic changes with destruction of tumour tissue.** | **(+2) Mild** |
| **5** | **DXY** | The tumour tissue comprised of variable sized cells of glands and fibroblasts as major cellular component. The tumour tissue comprised of variable sized cells of elongated and spindle shape with dense arrangement. Prominent basophilia with enlarged nuclei, mitotic figures and marked pleomorphism of fibroblastic neoplastic cells was noted.  **Multiple foci with showed marked necrotic changes with destruction of tumour tissue.** | **(+2) Mild** |
| **6** | **MET –DXY** | Moderately developed tumor tissue of proliferative fibrocytes and fibroblasts and glandular adenocarcinoma. The tumour tissue comprised of variable sized cells of glands and fibroblasts.  Moderate basophilia with enlarged nuclei, mitotic figures and marked pleomorphism of fibroblastic neoplastic cells was noted. The tumour tissue comprised of variable sized cells of elongated and spindle shape with dense arrangement.  **Significant and diffuse / multiple foci with degenerative and necrotic changes of neoplastic cells in the tumour mass.** | **(+3) Moderate** |
| **Note : Overall Grade score as- NAD** =No Abnormality Detected, **Minimal changes** (+1), **Mild changes** (+2), **Moderate changes** (+3), **Severe changes** (+4). | | | |

| **Sr. N.** | **Group /Slide Code** | B. **Histopathological observations**   **Kidney** | **Overall Pathological Grade** |
| --- | --- | --- | --- |
| **1** | **Control** | Normal histomorphology cellular features of renal tubules and glomeruli in the renal cortex and medulla region. Absence of inflammatory or pathological changes in renal parenchyma. | **NAD** |
| **2** | **MET** | Normal histomorphology cellular features of glomeruli in the renal cortex.  Focal and minimal degenerative changes of renal tubules.  Focal congested vascular tissue and interstitial hemorrhages in renal parenchyma.  Absence of inflammatory or pathological changes in renal parenchyma. | **Minimal (+1)** |
| **3** | **LEF** | Normal histomorphology cellular features of renal tubules and glomeruli in the renal cortex and medulla region. Absence of inflammatory or pathological changes in renal parenchyma. | **NAD** |
| **4** | **MET - LEF** | Normal histomorphology cellular features of renal tubules and glomeruli in the renal cortex and medulla region. Absence of inflammatory or pathological changes in renal parenchyma. | **NAD** |
| **5** | **DXY** | Normal histomorphology cellular features of renal tubules and glomeruli in the renal cortex and medulla region. Absence of inflammatory or pathological changes in renal parenchyma. | **NAD** |
| **6** | **MET –DXY** | Normal histomorphology cellular features of glomeruli in the renal cortex.  Focal and minimal degenerative changes of renal tubules.  Focal congested vascular tissue and interstitial hemorrhages in renal parenchyma.  Absence of inflammatory or pathological changes in renal parenchyma. | **Minimal (+1)** |
| **Note: Overall Grade score as- NAD** =No Abnormality Detected, **Minimal changes** (+1), **Mild changes** (+2), **Moderate changes** (+3), **Severe changes** (+4). | | | |

| **Sr. N.** | **Group /Slide Code** | **D. Histopathological observations**  **Lung** | **Overall Pathological Grade** |
| --- | --- | --- | --- |
| **1** | **Control** | Normal histomorphology features of bronchi and alveolar tissue with intact cellular details. Normal bronchiolar epithelium. | **NAD** |
| **2** | **MET** | Normal histomorphology features of bronchi and alveolar tissue with intact cellular details. Normal bronchiolar epithelium. | **NAD** |
| **3** | **LEF** | Focal areas of mononuclear cellular infiltration in the alveolar parenchyma.  Normal histomorphology features of bronchi and alveolar tissue with intact cellular details. | **Minimal (+1)** |
| **4** | **MET - LEF** | Normal histomorphology features of bronchi and alveolar tissue with intact cellular details. Normal bronchiolar epithelium. | **NAD** |
| **5** | **DXY** | Normal histomorphology features of bronchi and alveolar tissue with intact cellular details.  Normal bronchiolar epithelium. | **NAD** |
| **6** | **MET –DXY** | Focal areas of mononuclear cellular infiltration in the alveolar parenchyma.  Normal histomorphology features of bronchi and alveolar tissue with intact cellular details. | **Minimal (+1)** |
| **Note: Overall Grade score as-**  **NAD** =No Abnormality Detected, **Minimal changes** (+1), **Mild changes** (+2), **Moderate changes** (+3), **Severe changes** (+4). | | | |

| **Sr. N.** | **Group /Slide Code** | E. **Histopathological observations**   **Heart** | **Overall Pathological Grade** |
| --- | --- | --- | --- |
| **1** | **Control** | Normal histomorphology features of cardiac muscle fibers in the myocardium. Absence of inflammatory or pathological changes in heart tissue. | **NAD** |
| **2** | **MET** | Focal and minimal degenerative changes in the cardiac muscle fibers with focal areas of congestion and occasional foci of interstitial hemorrhages in pericardium and myocardium region. | **Minimal (+1)** |
| **3** | **LEF** | Normal histomorphology features of cardiac muscle fibers in the myocardium. Focal congested blood vascular tissue.  Absence of inflammatory or pathological changes in heart tissue. | **NAD** |
| **4** | **MET - LEF** | Normal histomorphology features of cardiac muscle fibers in the myocardium. Absence of inflammatory or pathological changes in heart tissue. | **NAD** |
| **5** | **DXY** | Normal histomorphology features of cardiac muscle fibers in the myocardium. Absence of inflammatory or pathological changes in heart tissue. | **NAD** |
| **6** | **MET –DXY** | Focal and minimal degenerative changes in the cardiac muscle fibers with focal areas of congestion and occasional foci of interstitial hemorrhages in pericardium and myocardium region. | **Minimal (+1)** |
| **Note: Overall Grade score as-**  **NAD** =No Abnormality Detected, **Minimal changes** (+1), **Mild changes** (+2), Moderate **changes** (+3), **Severe changes** (+4). | | | |

| **Sr. N.** | **Group /Slide Code** | **F. Histopathological observations**  **1. Liver** | **Overall Pathological Grade** |
| --- | --- | --- | --- |
| **1** | **Control** | Normal hepatic parenchyma with normal histomorphology of hepatocytes with intact cellular features. The hepatocytes were arranged in hepatic strands around central vein. Hepatocytes appeared polygonal to round in shape with presence of round nucleus and intact cell borders.  Absence of any metabolic or pathological cellular lesions in the liver. | **NAD** |
| **2** | **MET** | Focal areas of cellular swelling of hepatocytes.  Focal MNC infiltration in hepatic parenchyma. Congested central vein and portal veins. | **Minimal (+1)** |
| **3** | **LEF** | Focal areas of cellular swelling of hepatocytes.  Focal MNC infiltration in hepatic parenchyma. Congested central vein and portal veins. Focal vacuolar cytoplasmic changes in the hepatocytes. | **Minimal (+1)** |
| **4** | **MET - LEF** | Normal hepatic parenchyma with normal histomorphology of hepatocytes with intact cellular features. The hepatocytes were arranged in hepatic strands around central vein. Hepatocytes appeared polygonal to round in shape with presence of round nucleus and intact cell borders.  Focal congested central vein and portal veins.  Absence of any metabolic or pathological cellular lesions in the liver. | **NAD** |
| **5** | **DXY** | Mild pathological changes with areas of cellular swelling of hepatocytes with granular cytoplasmic changes and karyomegaly.  Congested central vein and portal veins.  Focal vacuolar cytoplasmic changes in the hepatocytes. | **Mild (+2)** |
| **6** | **MET –DXY** | Focal areas of cellular swelling of hepatocytes.  Focal MNC infiltration in hepatic parenchyma. Congested central vein and portal veins. | **Minimal (+1)** |
| **Note: Overall Grade score as- NAD** =No Abnormality Detected, Minimal **changes** (+1), Mild **changes** (+2), Moderate **changes** (+3), **Severe changes** (+4). | | | |
